# History-structured forecasting of rewarded give-up behavior in a rodent metacognition task

**DOI:** 10.64898/2026.08.11.744192

**Authors:** Bin Yin, Ya-Xin Wang, Chongyi Liu, Liya Fu

**Author notes:** These authors contributed equally (Bin Yin, Ya-Xin Wang).

## Abstract

Longitudinal animal experiments generate behavior that is individual, history-dependent, and sometimes affected by ordinary procedural irregularities, yet analyses commonly reduce such records to pooled averages or synchronous trial-level explanations. We introduce history-structured forecasting as an auditable framework for determining whether an animal’s own preceding behavior carries predictive information beyond current-trial context. We applied the framework to 213,990 events from rats performing an auditory duration-discrimination task, using leakage-safe chronological forward-chaining, explicit trivial baselines, and controls that reset, exchange, or disrupt behavioral history. A transparent gradient-boosted model achieved 51.7% four-class accuracy, exceeding last-action persistence (40.9%) and prefix-derived subject-modal prediction (37.2%); decline-class AUPRC was 0.525 against a prevalence baseline of 0.303. Validation showed that the predictive advantage depended predominantly on each animal’s short-range sequential action history rather than group-level history, subject identity alone, or reward/correctness features, and strengthened on genuine choice trials. Forecasting remained informative across all 23 labeled animals, including six with recoverable records affected by incorrect training programming. These results revise the interpretation of rewarded give-up behavior while demonstrating how recoverable irregular records can be retained in transparent robustness analyses. History-structured forecasting offers a reusable open-science strategy for extracting reproducible evidence from imperfect longitudinal animal records without creating an artificially clean cohort.

## Introduction

Metacognition in perceptual decision-making (specifically the decision to opt out, decline, or give up on a trial) is typically modeled as a synchronous, concurrent evaluation of evidence or confidence [1–8]. In dominant frameworks such as bounded accumulation and drift-diffusion models [9], reinforcement learning [10], and confidence models [11–15], give-up behavior occurs when momentary evidence for primary response alternatives fails to reach a critical threshold, or when uncertainty exceeds a static limit. In these accounts, the metacognitive decision is fundamentally a local calculation based on the immediate stimulus evidence and current task contingencies. Even within evidence- and value-based frameworks, choices are known to carry history-dependent biases such as choice hysteresis and gradual perseveration [16], but such effects are typically treated as nuisance deviations from an otherwise evidence-driven decision rather than as its primary determinant.

A useful framework distinguishes triggering causes from structuring causes, following Dretske’s [17] distinction between triggering and structuring causes of behavior, recently extended by Potter and Mitchell’s [18] discussion of causation and historicity in neuroscience. Triggering causes are proximal events that precipitate an action: the tone duration on the current trial, the availability of the lever or sensor, or the current reward ratio. Structuring causes are historically accumulated constraints that determine how such triggers are interpreted by the organism: learned action values, response habits, reward-history sensitivity, volatility of strategy, post-gap resetting, and context-specific policy stability. On this view, the same stimulus does not have a fixed behavioral meaning across subjects or across time. Its effect depends on the organized history that has shaped the animal’s current policy.

In the original experimental design, reward ratio was not a neutral background condition: later reward contexts were adjusted according to subjects’ prior decline tendencies. High-decline animals experienced increased reward for correct engagement, whereas low-decline animals experienced increased reward for the decline/contact-sensor option. Thus, reward ratio provided a key triggering context while also reflecting the subject’s prior behavioral history. Reward ratio was treated as a triggering cause at the current-trial level, but because reward-ratio blocks were assigned based on earlier individual decline tendencies, they also formed part of each subject’s diachronic learning environment.

This distinction motivates a forecasting framework that treats each subject’s behavior as a non-exchangeable longitudinal trajectory rather than a set of independent trials. If decline behavior is structured by accumulated behavioral history, then random trial-level train/test splits are inappropriate: they collapse the causal direction of learning history and allow future structure to inform past prediction.

We investigate an alternative diachronic hypothesis: that behavioral shifts in a metacognition task, particularly the decision to decline difficult trials, are heavily structured by an individual’s accumulated task history and sequential behavioral dynamics across sessions. Rather than a pure expression of momentary uncertainty, the give-up response functions as an iteratively learned, history-dependent behavioral strategy.

To rigorously test this hypothesis, we constructed an evaluation framework based on strict chronological forward-chaining. If decline behavior is fundamentally a temporally structured process rather than a static synchronous reaction, then predictive models incorporating subject-specific sequential behavioral history should improve on models restricted to local stimulus context; as we show, this predictive structure is carried predominantly by short-range, within-session sequential history rather than by a demonstrated slow cross-session consolidation. By requiring models to forecast future behavior on entirely unseen late-stage sessions, we rule out retrospective data fitting and ensure that the identified structures represent out-of-sample predictive signals. In an exploratory analysis, we also show that free-running closed-loop simulations can exhibit an over-decline feedback tendency, highlighting a boundary between next-step forecasting and stable behavioral generation rather than serving as primary evidence for the structuring-cause claim.

## Results

### Dataset: cross-subject heterogeneity

We reconstructed 24 dataset-level subject trajectories from rats performing an auditory duration-discrimination task, spanning 213,990 trial events across 2,162 sessions; the task apparatus, trial contingencies, and reconstruction pipeline are summarized in Figure 1. Forecasting requires the trial-level action as its target, and 23 of the 24 trajectories carry these labels; because our approach is individual and history-structured, we modeled all 23, with no labeled subject excluded on behavioral or performance grounds. The 17 subjects with both raw event logs and derived trial-level records provide the primary held-out evaluation. Six separate raw-only subjects lacked the derived trial-level tables used for core validation and had initially been set aside after periods of incorrect training programming; because their histories and action labels were recoverable, they enter the all-23 robustness analysis (below). The remaining dataset-level trajectory, RED 4, is not one of these six and has no action labels in any of its 668 phase-coded late-discrimination rows, and therefore no forecasting target; we note it here for completeness but do not include it in modeling (Methods). The dataset revealed significant behavioral heterogeneity. Trials occurred across reward-ratio contexts that modulated the relative value of lever-based correct engagement and contact-sensor decline. Base decline rates ranged widely across individuals, from less than 5% to over 60%. Furthermore, decline actions exhibited distinct reaction time (RT) profiles compared to standard engagement or omission actions, indicating that giving up is a distinct, non-random action rather than a byproduct of simple distraction or failure to act. Under the current reward-ratio context, validated decline actions were rewarded, whereas incorrect engagements or omissions yielded no reward and triggered a timeout.

**Figure 1.**
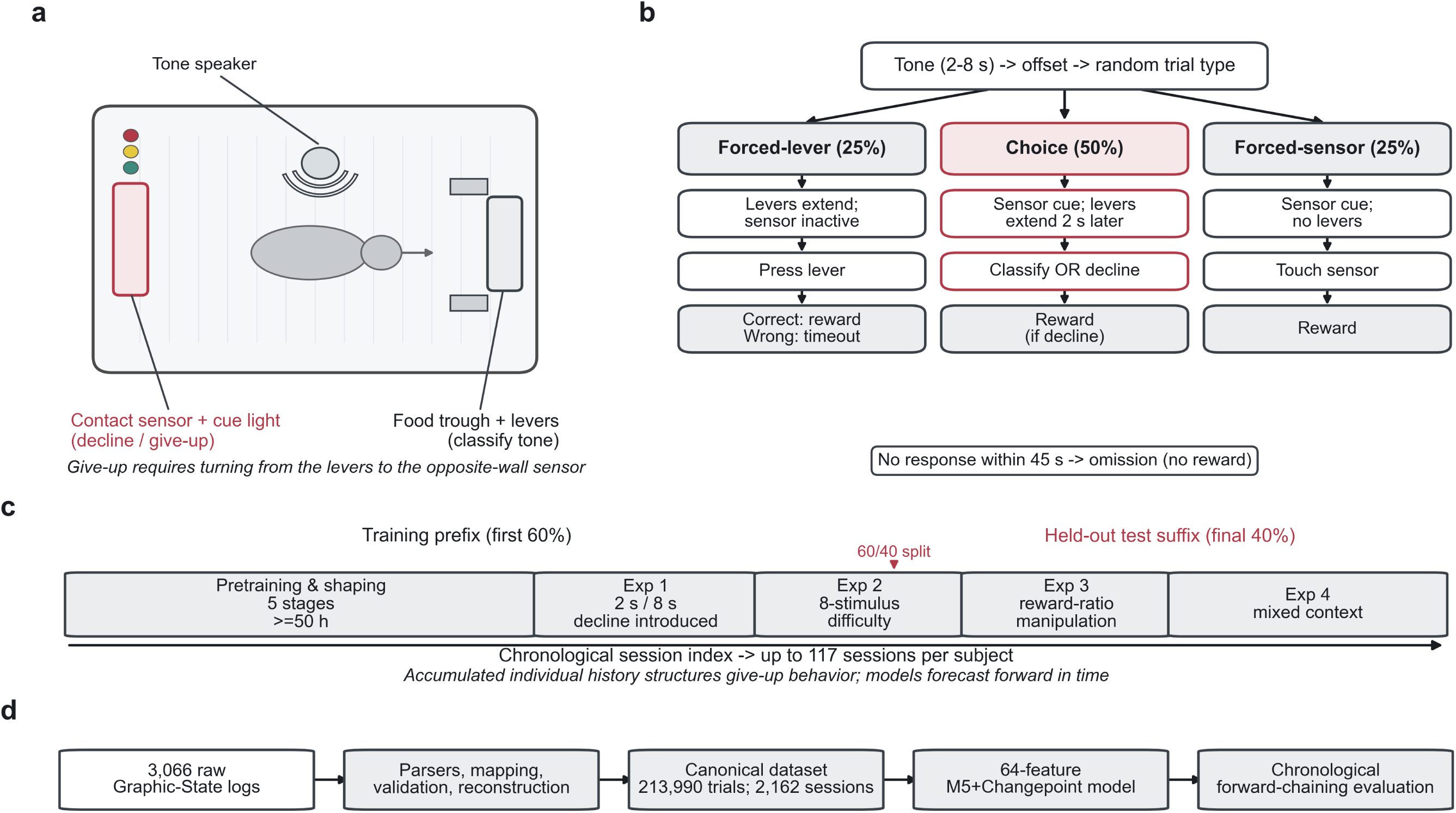
Task, learning history, and analysis pipeline. (a) Top-down schematic of the operant chamber: the contact sensor with its tri-colour cue light (the rewarded decline/give-up option) sits on the left wall, opposite the food trough and left/right classification levers, with the tone speaker overhead; giving up requires turning away from the levers to the sensor. (b) The three post-tone trial types: forced-lever (25%), choice (50%; classify by a lever press or decline by touching the sensor), and forced-sensor (25%); rewards, timeouts and the 45-s omission window follow the current reward-ratio context (Methods). (c) Diachronic learning history across four phases (Exp 1-4: decline introduction, eight-stimulus difficulty, reward-ratio manipulation, mixed context) over up to 117 sessions; the dashed line marks an illustrative per-subject 60/40 chronological split (Supplementary Figure S3). (d) Pipeline: 3,066 raw Graphic-State logs → parsing, mapping, validation and reconstruction → the canonical dataset (213,990 trials, 2,162 sessions) → the 64-feature M5+Changepoint model → chronological forward-chaining evaluation.

### Individual diachronic history supports the structuring-cause account

To test whether individual diachronic history provides predictive value beyond group statistics and synchronic subject identity, we compared eight models spanning a 2×2 taxonomy (group vs individual × synchronic vs diachronic) plus three temporal-order controls (Figure 2a).

**Figure 2.**
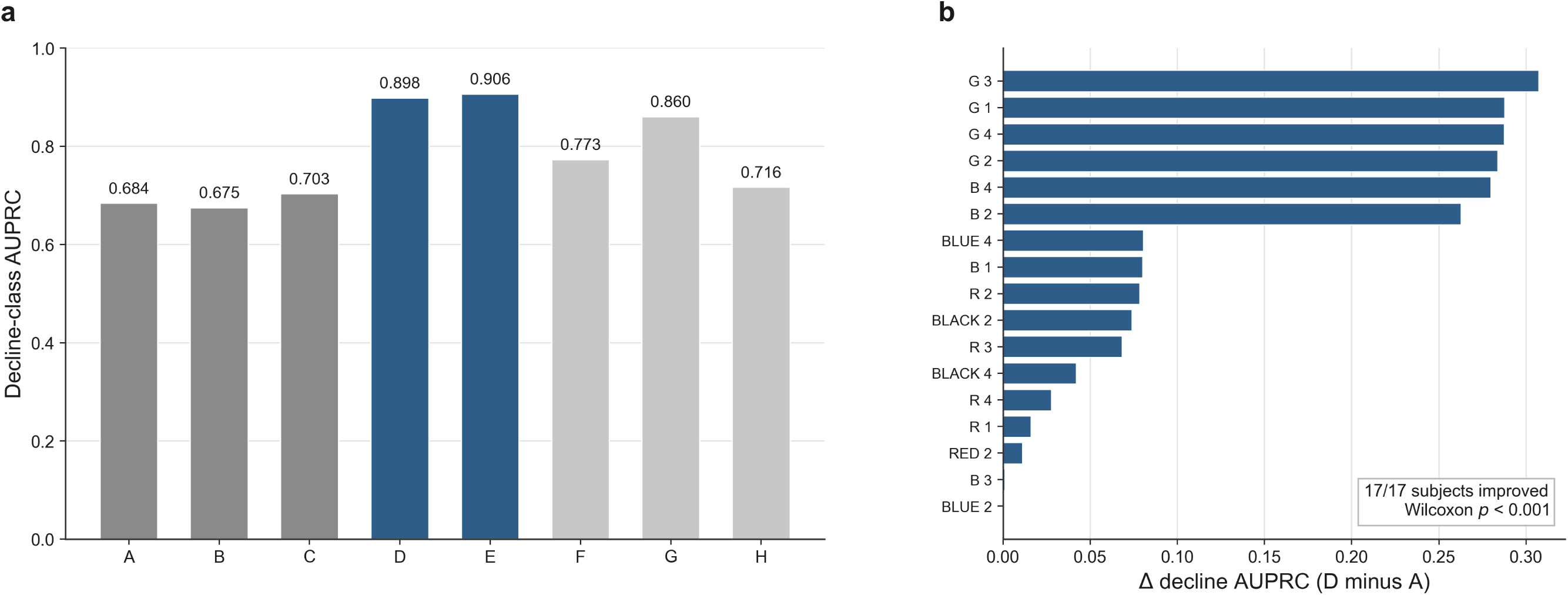
Individual diachronic history carries the dominant structuring signal for decline behavior. (a) Decline-class AUPRC for the eight-model taxonomy: triggering-only (A), group-history (B), subject-static (C), individual-history (D), full M5+Changepoint (E), time-scrambled (F), session-shuffled (G) and cross-subject history swap (H). The focal individual-history models D and E are shown in blue; comparison and control models are shown in grey. (b) Per-subject advantage in decline AUPRC of individual-history D over triggering-only A, ordered by magnitude; all 17 subjects improved (paired mean +0.129, bootstrap 95% CI [0.074, 0.186], Wilcoxon signed-rank *p* < 0.001).

Group-level history statistics showed no credible evidence of predictive value beyond current triggering context (Model B aggregate accuracy 52.4% versus Model A 55.4%, and decline AUPRC 0.675 versus 0.684). Subject identity without trial-level history improved accuracy by 9.4 percentage points (Model C) but degraded probability calibration (log loss 1.147 vs 1.060). Individual trial-by-trial history (Model D) improved aggregate performance over triggering-only (Model A): accuracy 66.8% vs 55.4%, decline AUPRC 0.898 vs 0.684, and log loss 0.894 vs 1.060. At the subject-paired level, D improved decline AUPRC over A by +0.129 on average, with all 17 of 17 subjects improving (Figure 2b; Supplementary Figure S7). In the group-history contrast (Figure 2a, models B versus D), D improved over B by paired mean +0.137, Wilcoxon *p* < 0.001 (17 of 17 subjects; per-subject values in Supplementary Data 5). Changepoint features added a small increment to decline detection that did not reach significance (E vs D aggregate AUPRC: 0.906 vs 0.898; paired Wilcoxon *p* = 0.08). The individual-history features therefore encoded not only prior choices and rewards, but also the animal’s realized trajectory through reward-ratio contexts.

Three temporal-order controls confirmed that the predictive signal depends on chronological, subject-specific sequential structure. Because these controls alter history depth as well as order, we compared them against a depth-matched chronological model (D with history recomputed within the same evaluation blocks); this depth control did not itself change decline AUPRC (D vs depth-matched D, +0.0003, *p* = 0.28). Time-scrambling individual history (Model F) degraded aggregate decline AUPRC from the depth-matched chronological anchor D(reset) (0.906) to 0.773 (paired mean difference +0.118, Wilcoxon *p* < 0.001), demonstrating that the temporal ordering of experience, not merely its marginal statistics, carries the structuring signal (Supplementary Figure S8). Substituting another subject’s history (Model H) similarly degraded performance (aggregate AUPRC 0.716; paired difference +0.099, *p* < 0.001), confirming that the signal is individual. Session-shuffling (Model G), by contrast, preserved accuracy and decline AUPRC (paired difference +0.008, *p* = 0.07), with only a borderline effect on log loss (*p* = 0.07): the order of whole sessions matters little, whereas the fine-grained trial order and the animal’s ownership of its history do. Pairwise subject-level comparisons were treated as planned contrasts within the structuring-cause validation family; the primary D-vs-A and D-vs-B conclusions remain significant under conservative Bonferroni correction across the eight reported model contrasts (all uncorrected *p* < 0.001; corrected *p* < 0.001).

Absolute AUPRC and accuracy values in this validation analysis exceed the primary M5+Changepoint benchmark (decline AUPRC 0.525, accuracy 51.7%) because, unlike the headline model, every model here includes the current trial type as a covariate. Because forced-sensor trials make a decline near-mandatory, trial type inflates the absolute decline AUPRC of all eight models alike (triggering-only Model A already reaches 0.684); accordingly all conclusions rest on within-analysis paired contrasts, not absolute values. To confirm the individual-history advantage is not an artifact of mandated or otherwise deterministic trial types, we recomputed the D-vs-A contrast on genuine choice trials only (25,036 trials, decline base rate 0.21): the advantage grew rather than shrank, to a per-subject mean of +0.253 decline AUPRC (16 of 17 subjects, exact Wilcoxon *p* = 4.6 × 10^-5; pooled Model A 0.208 versus Model D 0.671). Because this restriction removes all non-choice categories rather than forced-sensor trials alone, it shows that deterministic non-choice trials do not manufacture the individual-history signal and collectively dilute its within-choice contrast (Supplementary Information S9).

### History-based models improve on static context

The full model comparison is summarized in Figure 3a–c and Supplementary Data 2. When predicting trial-by-trial behavior (a 4-class outcome: left, right, decline, omit; chance 25%), models restricted to static trial context, including stimulus and reward-ratio context variables, underperformed substantially relative to history-based models. A stimulus-aware logistic regression (M1) evaluated on a strict 60/40 chronological split achieved only 42.9% accuracy, a log loss of 1.587, and a decline-class AUPRC of 0.481.

**Figure 3.**
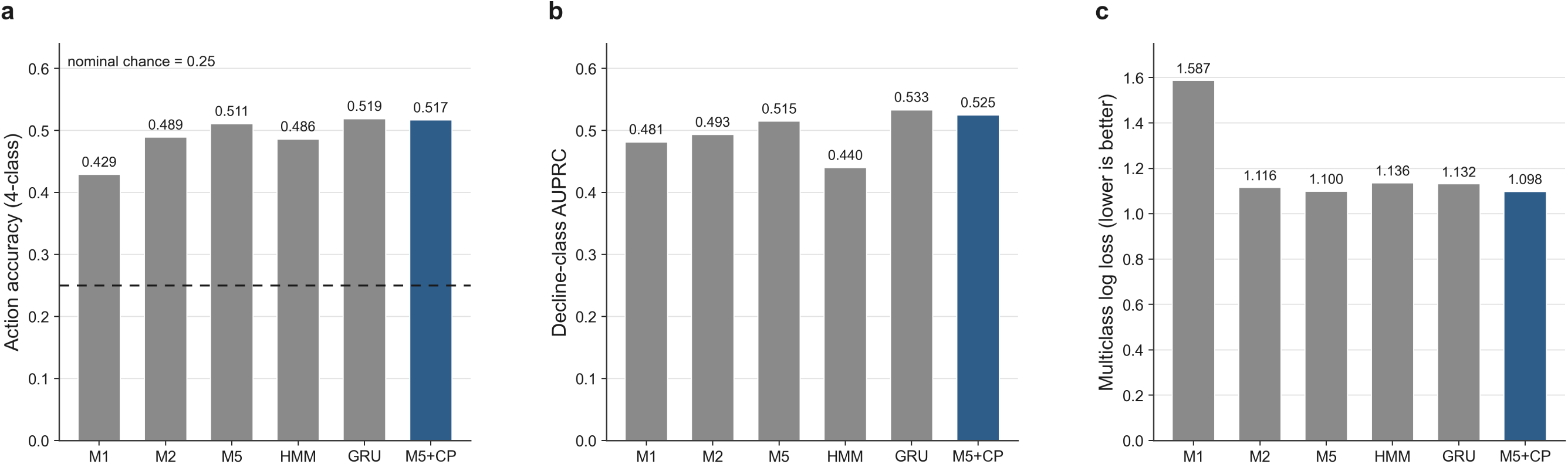
Comparison of a transparent history-based model with static, latent and recurrent baselines. All models used chronological prefix-to- suffix splitting. The fresh temperature-scaled M5+Changepoint result is shown in blue; archival screening comparators are grey. (a) Four-class action accuracy (dashed line, nominal chance 25%). (b) Decline-class AUPRC (the decline base rate is 0.30 in this test set, so 0.525 is a lift over prevalence). (c) Multiclass log loss (lower is better). M1, stimulus-aware logistic regression; M2, sequential LightGBM; M5, EWM+history LightGBM; HMM, *K* = 4 hidden Markov model; GRU, gated recurrent unit (*h* = 8); M5+Changepoint, M5 with online changepoint features. The eligible-trial sets differ across model families (e.g., 39,237 trials for GRU versus 51,090 for M5+Changepoint), so the comparison supports model-family screening rather than exact paired superiority.

In contrast, sequential baselines performed substantially better. Incorporating lagged actions, running rates, and exponentially weighted moving (EWM) averages provided a large forecasting improvement. The EWM+history LightGBM model (M5: lagged actions, running rates and EWM averages) [19] achieved 51.1% accuracy, a log loss of 1.100, and a decline AUPRC of 0.515 (the plain sequential LightGBM, M2 in Figure 3, is a weaker screening comparator). In the archival Phase-3 screening, static aggregated early-phase summaries showed no credible evidence of improving prediction beyond subject identity and recent sequential dynamics (Supplementary Data 2); this secondary comparison was not rerun on the canonical stack. This does not rule out that early experience contributes through pathways not captured by those summary statistics. An archival feature-group ablation is reported descriptively in Supplementary Figure S9.

### Online changepoint features and the final model

Behavioral strategy is non-stationary across sessions. An archival PELT [20] segmentation of session-level statistics identified 131 distinct strategy shifts across 23 subjects; this descriptive characterization motivated, but is not itself part of, the released feature pipeline. The released M5+Changepoint model instead computes online change-sensitive proxies causally from each subject’s prefix (short- versus long-span EWM deltas, action-run age, and a post-gap flag), added to the EWM LightGBM model. This model achieved a calibrated log loss of 1.098, an action accuracy of 51.7%, and a decline AUPRC of 0.525 (Figure 3a–c). Adding the online changepoint features to the individual-history model gave only a small, non-significant increment over history alone (E vs D: +0.6 pp accuracy, +0.010 decline AUPRC, *p* = 0.080), so the changepoint component refines rather than drives the forecast.

### Recurrent and latent model comparisons

We evaluated deep sequence models (GRU with *h* = 4 and *h* = 8) [21] to capture non-linear temporal integration. On their smaller eligible-trial set the best GRU matched or marginally exceeded the transparent LightGBM model on accuracy and decline AUPRC; we retain the transparent M5+Changepoint model for its interpretability, calibration and matched evaluation index rather than for superior discrimination. The best GRU configuration achieved a log loss of 1.132, close to M5+Changepoint’s 1.098, and a decline AUPRC of 0.533 (Figure 3b,c; 39,237 trials, versus 0.525 on 51,090 for M5+Changepoint). GRUs also yielded substantially worse RT prediction (MAE > 7 s). Model comparisons across phases should be interpreted as stage-specific evaluations under their documented eligible-trial sets; the final M5+Changepoint benchmark and the structuring-cause validation use fixed evaluation indices. Where evaluation sets differ (e.g., GRU models were evaluated on 39,237 trials vs. 51,090 for M5+Changepoint), comparisons support model-family screening rather than exact paired superiority claims.

Similarly, we fit a categorical *K* = 4 hidden Markov model (HMM) [22] to test if latent states capture structure better than transparent history. The HMM segmented behavior into four empirical clusters (Disengaged, Decline-heavy, Left-engaged, Right-engaged), but the Disengaged state proved to be near-absorbing (mean dwell 928 trials). A properly forward-chained HMM achieved only 48.6% accuracy and an AUPRC of 0.440, falling well below the transparent M5 floor (Figure 3a,b; Supplementary Figure S5). The sequential structure of give-up/decline behavior is thus effectively captured by transparent moving averages and changepoint flags.

### The final teacher-forced model

The final predictive architecture (M5+Changepoint), summarized in Figure 4, consolidates these findings into two components:

1. **Action prediction.** A temperature-scaled M5+Changepoint LightGBM [19,23] (*T* = 1.347) using 64 features. On 51,090 held-out test trials, it achieved 51.7% accuracy, a log loss of 1.098, a decline AUPRC of 0.525, a decline F1 of 0.424, and an expected calibration error (ECE) of 0.036 (Brier 0.180; Figure 4a,c). On the same 51,090-trial test set this accuracy exceeds trivial non-model baselines: last-action persistence (40.9%; the model is better in 15 of 17 subjects), each subject’s prefix-derived modal action (37.2%; 17 of 17) and a conservative held-out modal rate (40.2%; 17 of 17); the decline AUPRC of 0.525 is +0.22 over the 0.30 decline prevalence, and 16 of 17 subjects exceed their own decline-prevalence baseline (Supplementary Data 8; baselines from scripts/14).
2. **Dedicated reaction-time modeling.** An action-conditioned lognormal LightGBM architecture substantially improved reaction-time prediction. The dedicated RT model achieved a standalone mean absolute error (MAE) of 1.43 s, which improved to 1.26 s in the integrated simulation pipeline, a 32–40% improvement over generic shared-backbone architectures (Figure 4b; Supplementary Figure S6).

**Figure 4.**
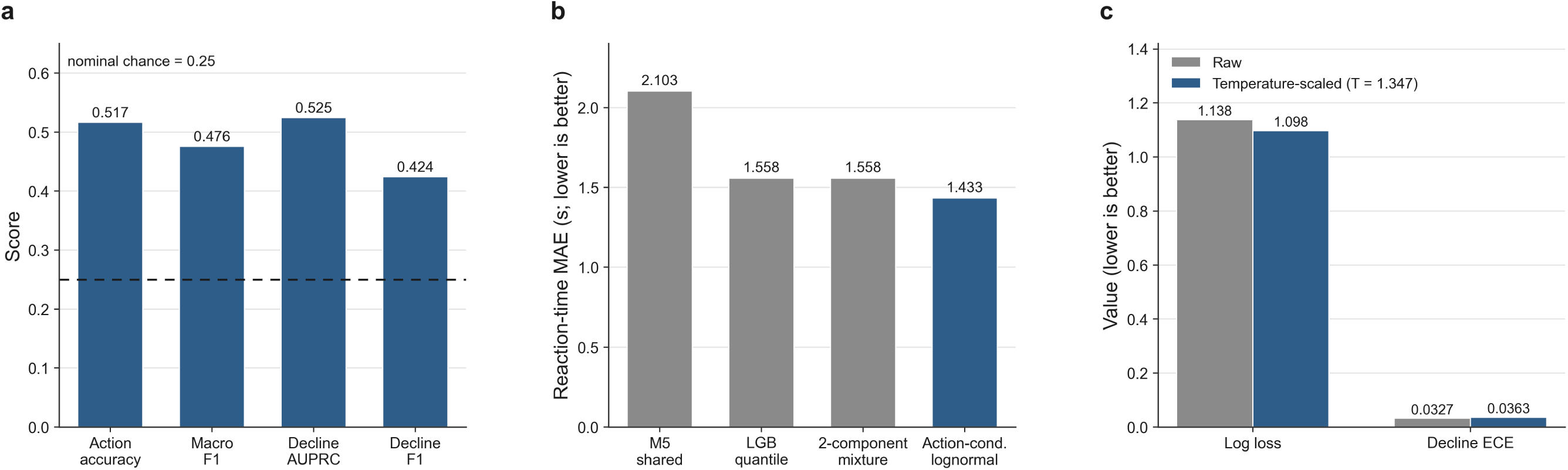
Teacher-forced model performance. (a) Headline metrics of the temperature-scaled M5+Changepoint model on 51,090 held-out test trials (dashed line, nominal chance 25%). (b) Archival reaction-time model comparison (mean absolute error; lower is better): the action-conditioned lognormal model is compared with the shared-backbone M5 baseline, a quantile model and a two-component mixture. (c) Fresh calibration comparison. Temperature scaling (*T* = 1.347) reduced multiclass log loss from 1.138 to 1.098 while decline ECE changed from 0.0327 to 0.0363; lower values are better for both metrics.

### Individual structuring-state profiles

Beyond aggregate performance, the non-tautological structuring-state features extracted from the calibrated model (Methods) summarize each subject’s realized reward-sensitive tendencies across its chronological trajectory. These profiles varied markedly across individuals, most conspicuously in win-stay rate, indicating that the predictive signal is carried by individualized, history-accumulated policy states rather than a strategy shared across the cohort (Figure 5; Supplementary Data 3).

**Figure 5.**
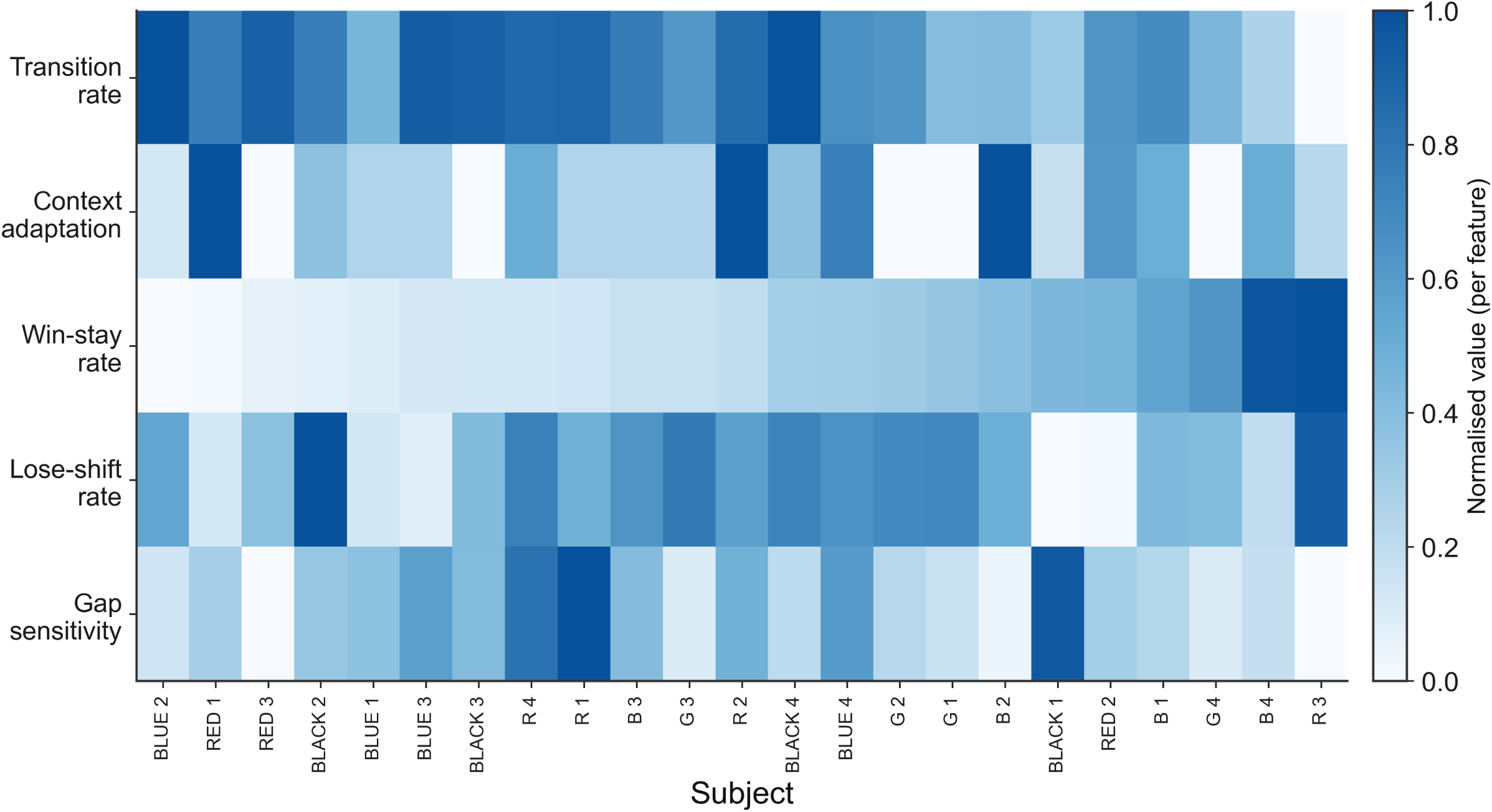
**Individual structuring-state (***c_t_***) profiles.** Heatmap of five non-tautological *c_t_* features (transition rate, context adaptation, win-stay rate, lose-shift rate and gap sensitivity), min-max normalised per feature across 23 subjects, showing distinct individual structuring strategies. The *c_t_* feature definitions follow the archival Phase-3 derivation (simplified relative to the fresh benchmark pipeline) and are provided as descriptive per- subject profiles (Supplementary Data 3). (Free-running simulation reliability, an exploratory model-boundary diagnostic, is reported in Supplementary Information S7 rather than as a main-figure panel.)

### Robustness across all labeled subjects, including procedure-disrupted animals

To confirm that the primary result is not specific to the 17 core-validated subjects, we repeated the forecast on all 23 labeled subjects under an identical leakage-safe chronological split (each subject’s first 60% of sessions for training, its final 40% held out; RED 4 is not modeled, having no action labels). This is a within-subject chronological forecast across all 23 animals. Pooled action accuracy was 52.0%, compared with 41.0% for last-action persistence, 35.4% for each subject’s prefix-derived modal action, and 41.7% for a conservative held-out modal rate; the model beat persistence in 21 of 23 subjects, the deployable prefix-modal predictor in 23 of 23, and the conservative held-out modal rate in 18 of 23 (Figure 6a; Supplementary Data 8). Calibrated decline AUPRC was 0.514, +0.219 over the 0.295 decline prevalence, and 22 of 23 subjects exceeded their own decline-prevalence baseline (Figure 6b; the exception was BLUE 2). The 17-core subset reproduced the primary values to reported precision (51.7%, 0.525). Per-subject accuracy ranged from 43% to 66%. The 6 raw-only subjects, all initially set aside after periods of incorrect training programming, were forecast on their held-out final 40% comparably to the core cohort (55.7% accuracy), showing that within-subject forecasting remains viable in the procedurally irregular records. An exploratory parameter-transfer analysis for these six animals (models fitted only to the 17 core animals, while using each target animal’s available prior history) is reported in the Supplementary Information (Supplementary Data 6, Supplementary Figure S10a–c). Together, these analyses show that above-nominal within-subject forecasting extends across the procedurally heterogeneous records; they do not establish broad cross-subject transfer.

**Figure 6.**
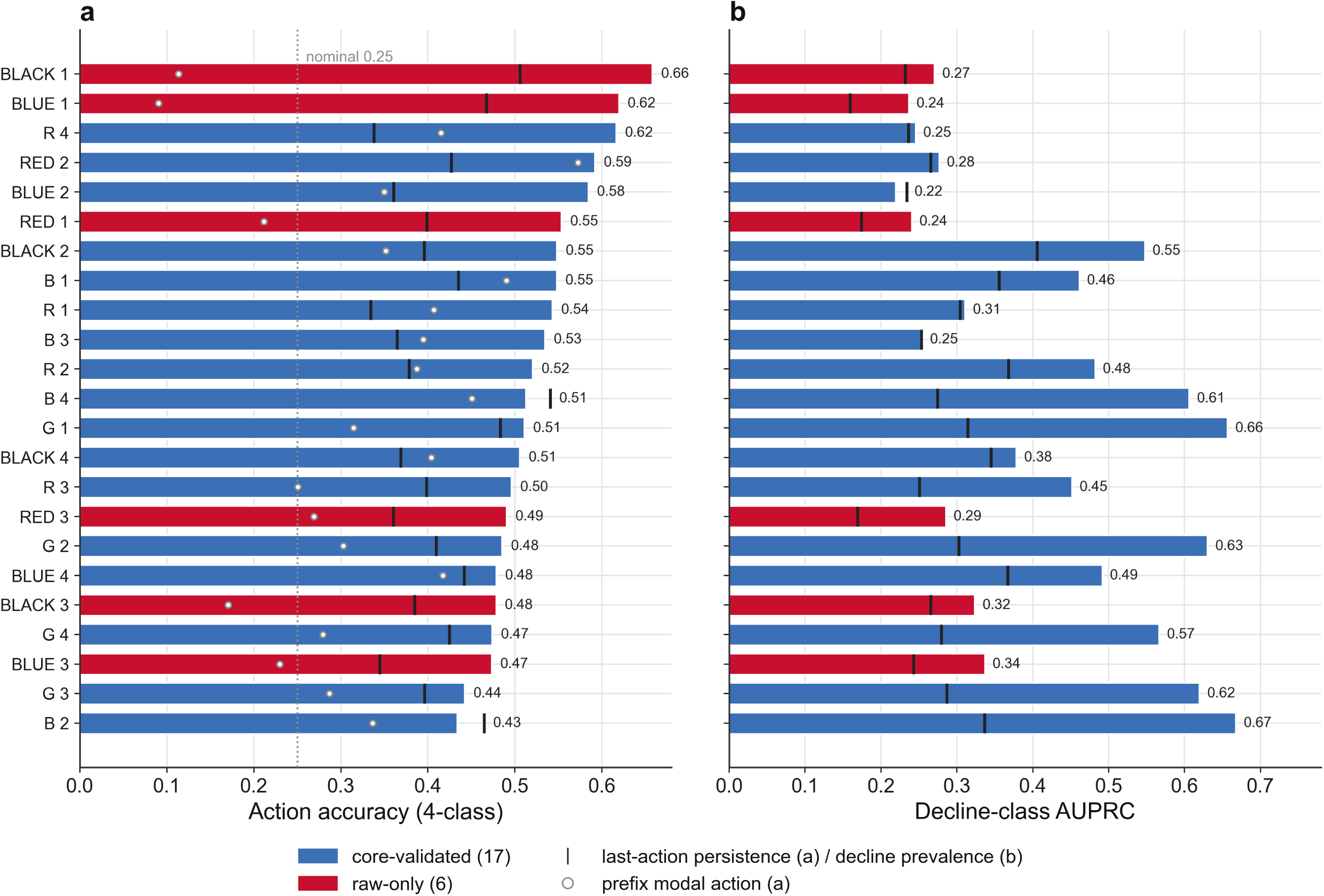
Per-subject forecasting across all 23 labeled subjects, benchmarked against trivial baselines. The M5+Changepoint model was evaluated under an identical leakage-safe chronological split on every labeled subject (RED 4 excluded, having no action labels). (a) Per-subject four-class action accuracy (bars) with each subject’s last-action persistence (black tick) and prefix-derived modal action (grey circle); the dotted line marks the nominal 0.25 four-class rate as a visual reference, not the inferential baseline. Bars are coloured 17 core-validated (blue) and 6 raw-only (red; initially set aside after periods of incorrect training programming). Pooled, the model beats last-action persistence in 21 of 23 subjects, the prefix-derived modal action in 23 of 23, and a conservative held-out modal rate in 18 of 23. (b) Per-subject decline-class AUPRC (bars), same subject order, with each subject’s decline prevalence (black tick); AUPRC is read as lift over prevalence, and 22 of 23 subjects exceed their own decline-prevalence baseline (the exception is BLUE 2). Pooled accuracy is 52.0% and calibrated decline AUPRC 0.514, +0.219 over prevalence (Supplementary Data 8), close to the 17-core primary and confirming the result is not specific to the most complete records.

### Exploratory closed-loop simulation suggests a generative boundary

To evaluate the boundary between step-ahead forecasting and behavioral generation, we also examined free-running closed-loop simulations in which model predictions were autoregressively fed back into subsequent history features (Supplementary Figure S12a–d). Because the exact magnitude of the closed-loop over-decline bias is sensitive to simulation implementation details, we treat this analysis as exploratory. It is not used as primary evidence for the structuring-cause claim. The theoretical point is narrower: teacher-forced prediction is not the same as closed-loop behavioral generation. A model that forecasts the next action from observed chronological history can drift when its own rewarded give-up predictions become part of the future history it must condition on.

## Discussion

These findings emphasize that give-up/decline behavior in this rodent metacognition task is not explained by momentary stimulus uncertainty alone. The central contribution is that each subject’s own chronological learning history, rather than group history or temporally exchangeable summaries, provides the main structuring signal for forecasting future rewarded give-up behavior. The ability to predict a give-up response with 51.7% accuracy, well above last-action persistence and each subject’s modal action, based on sequential and temporal features demonstrates that putative metacognitive strategy in this paradigm is strongly shaped by subject-specific sequential behavioral history (predominantly action history rather than an independently isolable reinforcement-history contribution). We therefore interpret give-up choices as putative metacognitive responses embedded in an operant reward structure, rather than as direct reports of confidence.

This diachronic account stands in productive tension with the dominant synchronic framing of opt-out behavior. In influential confidence-based models, the decision to decline or seek an alternative is read out from a momentary estimate of decision confidence computed from the current stimulus and its accumulated evidence [2,5,11–13], with candidate neural correlates identified in orbitofrontal and parietal circuits [6–7]. Consistent with an integrative rather than purely synchronic view, recent single-neuron work in a macaque opt-out task shows that metacognitive decline decisions are predicted by the integration of momentary working-memory strength with trial history and pre-stimulus arousal [24], indicating that accumulated history and current evidence jointly shape opt-out rather than the former simply replacing the latter. Such accounts capture graded opt-out as a function of trial difficulty and have been highly generative. Our results do not deny a role for momentary uncertainty, but they show that over the extended timescale of an operant metacognition paradigm the same nominal difficulty does not map onto a fixed decline probability: its behavioral meaning is set by the sequential behavioral history that has shaped each animal’s current policy. Where confidence-readout models typically treat the mapping from evidence to opt-out as given, the triggering-versus-structuring-cause framework [17–18] makes the learning and consolidation of that mapping the primary explanatory target, and the forward-chained forecasting result quantifies how much of future behavior this accumulated structure determines.

This distinction bears directly on the comparative study of metacognition. Demonstrations of uncertainty monitoring in rats and other non-human animals have often hinged on whether opt-out or decline responses track objective difficulty in the predicted direction [1,3,8,14–15], a synchronic criterion evaluated within or across sessions. Our finding that the give-up response is heavily structured by individual sequential behavioral history urges caution in reading such responses as direct reports of a metacognitive state: a rewarded decline option embedded in a reward schedule can come to be governed by learned action values and reward-history sensitivity, yielding difficulty-tracking behavior that need not index online confidence, consistent with a long-standing debate over associative versus metacognitive explanations of apparent metacognition in non-human animals [25–26]. Rather than adjudicating for or against animal metacognition, longitudinal, history-aware modeling offers a principled way to separate a genuinely synchronic confidence signal from a diachronically learned policy, a separation that single-session, cohort-aggregated analyses cannot make.

Because reward-ratio blocks were selected according to prior decline tendency, the external task environment was itself partially shaped by earlier individual behavior. This design makes the dataset especially appropriate for structuring-cause analysis: later behavior is not predicted from a static context alone, but from the interaction between current reward contingencies and each subject’s accumulated history. Reward ratio should therefore be interpreted as an important triggering context and historically contingent environmental condition, not as the only cause of decline behavior. The rewarded give-up response can also be viewed, by analogy, as a patch-leaving decision: recent foraging work models the leave/stay choice as a competitive integration of elapsed time and accumulated reward against a slowly varying latent state [27]. We draw this only as an analogy rather than an identification of the present task with foraging, but it likewise separates an economic disengagement policy from a synchronic confidence readout.

Teacher-forced prediction is not the same as closed-loop behavioral generation. The exploratory closed-loop simulation highlights the limits of phenomenological predictive modeling: transparent history features can support single-step prediction, yet drift when model-generated actions and rewards are recursively fed back as future inputs. The closed-loop simulation result should be interpreted as a model-boundary analysis rather than as evidence that real animals or the final public model exhibit a fixed attractor of this exact magnitude.

The structuring-cause validation analysis provides converging evidence that trial-by-trial decline behavior is shaped by each subject’s accumulated individual history in a way that is sequential, temporally ordered, and subject-specific. Group-level statistics showed no credible evidence of adding predictive information beyond current stimulus context, and subject identity without dynamic history was insufficient for probability calibration. These findings are consistent with the distinction between triggering and structuring causes (Dretske [17]; Potter and Mitchell [18]): the same stimulus does not have a fixed behavioral meaning across subjects or across time; its effect depends on the organized history that has shaped each animal’s current decision policy.

Within-subject forecasting beat last-action persistence and each subject’s prefix-derived modal action for all six raw-only subjects (though it beat the conservative held-out modal rate in only one of six), indicating that the method can accommodate heterogeneous records when each animal’s earlier history is available. The separate six-animal parameter-transfer probe was exploratory and did not establish an individual-history advantage or broad cross-subject generalization (Supplementary Information S10). A practical consequence follows from conditioning on each subject’s own accumulated history rather than restricting analysis to clean, protocol-conforming records: sessions affected by procedural irregularities may still contain usable longitudinal information. The model beat each of the 23 labeled subjects’ prefix-derived modal action (23 of 23) and last-action persistence in 21 of 23, including animals with documented training irregularities (Figure 6). History-structured forecasting is therefore applicable to procedurally heterogeneous longitudinal data, although its transfer to wholly new subjects or paradigms requires dedicated validation.

The temporal-order controls further sharpen this conclusion. Destroying chronological order within subjects substantially reduced decline prediction (decline AUPRC from 0.906 to 0.773) despite preserving action-rate marginals, demonstrating that the sequence of experience, not merely its summary statistics, carries the structuring signal. The partial preservation of signal under session-shuffling indicates that within-session dynamics (trial-to-trial transitions and running rates) account for most of the discriminative signal, with whole-session order contributing comparatively little; the cross-session contribution is small and at best borderline (a decline-AUPRC difference of +0.008, borderline by the paired test; Supplementary Information S9). The predictive signal is therefore predominantly short-range, within-session sequential behavioral history, rather than a demonstrated slow cross-session consolidation of decision criteria.

The individual structuring-state profiles (*c_t_*; Figure 5) may further help explain why some subjects were more vulnerable to drift in the exploratory closed-loop diagnostics: subjects with the most reward-sensitive win-stay tendencies are precisely those whose future history is most reshaped when their own rewarded give-up predictions are fed back as inputs (Supplementary Information S7). Models of give-up behavior in animal metacognition tasks should explicitly account for chronological strategy drift. Static snapshot analyses risk misinterpreting learned policy artifacts as immediate confidence readouts. Our reliance on interpretable, per-subject history features is complementary to a recent movement toward transparent behavioral models, including tiny recurrent networks that recover parsimonious accounts of individual reward-learning choices [28], and to evidence that human reward learning is shaped by memory variables beyond incremental value updating [29]; relative to more opaque predictors, our contribution is to make subject-specific sequential behavioral history the explicit, auditable signal. Future work simulating long-term behavioral trajectories will likely require meta-reinforcement learning or active inference architectures capable of representing and updating latent belief states to break the feedback tendencies of autoregressive historical models.

### Generalizability to longitudinal behavioral paradigms

The framework’s reach extends well beyond the present paradigm. Although this analysis concerns rewarded give-up behavior in a rodent metacognition task, the methodological logic is not task-specific. Many paradigms in psychology and behavioral neuroscience generate non-exchangeable individual trajectories in which current choices are shaped by prior feedback, reward exposure, errors, omissions, confidence states, and strategy shifts. The same framework (preserving each subject’s chronological order, engineering interpretable history features, and forecasting future behavior under leakage-safe forward-chaining while comparing individual diachronic history against current context, static subject profiles, group-level history, and temporally disrupted controls) can be applied to human and animal paradigms involving opt-out decisions, confidence judgments, reinforcement-learning and bandit tasks, effort allocation, self-regulated learning, emotion-regulation choice, and digital-phenotyping or intervention-engagement data. The requirement is not a particular species or apparatus but a longitudinal decision structure with repeated feedback. Importantly, what transfers is the analysis strategy rather than the specific result: establishing that individual chronological history carries a structuring signal in any new paradigm requires re-running the same structural controls there, not assuming the rodent finding generalizes.

### Limitations

Several limitations qualify the interpretation of these findings and indicate directions for future work.

1. *Closed-loop simulation is exploratory.* The free-running simulation is consistent with a reward-history feedback mechanism, but we retain it as exploratory: its over-decline magnitude is configuration-dependent rather than a single value (see Supplementary Information S7), and its per-subject generative reliability is low (Supplementary Information S7). It is not used as primary evidence and should not be relied upon as a synthetic-subject generator for arbitrary closed-loop testing.
2. *Early-history causality.* Static aggregated early-phase summaries showed no credible evidence of improving prediction beyond subject identity and recent sequential dynamics, though this does not rule out that early experience contributes through pathways not captured by these summary statistics.
3. *HMM ontology.* The identified HMM states (e.g., “Left-engaged”, “Disengaged”) are purely empirical statistical clusters of sequential choice behavior, lacking external physiological or subjective psychological grounding.
4. *Sex as a biological variable.* All subjects were male Sprague-Dawley rats; sex-specific generalization was not tested.

## Methods

### Operationalizing triggering vs. structuring causes

Following the triggering/structuring-cause framework (Dretske [17]; Potter and Mitchell [18]), we operationalize the causal inputs to our forecasting models into two classes. Triggering causes are defined as current-trial stimulus and context variables (e.g., the true stimulus duration, reward-ratio context, the immediate task contingencies). Structuring causes are defined as history-dependent variables derived from the causal prefix of the subject’s experience. These include lagged actions, exponentially weighted moving (EWM) averages of behavioral rates, cumulative reward histories, inter-session gap durations, and local strategy changepoint flags. This separation allows us to rigorously compare the predictive power of momentary triggers against accumulated structural constraints.

### Structuring-cause validation protocol

To test whether individual diachronic history carries structuring-cause-like predictive information for trial-level behavior, we trained eight LightGBM multiclass models spanning a 2×2 taxonomy (group-level vs individual-level × synchronic vs diachronic) plus three temporal-order controls. All models shared the same 51,090-trial test set, hyperparameters (num_leaves = 63, min_child_samples = 50, learning rate = 0.05, class_weight = balanced), and temperature-scaling calibration protocol. The triggering-only model (A) used only current-trial variables (stimulus duration, stimulus class, difficulty, trial type, paradigm, context). The group-history model (B) added group-level action-rate statistics from the training prefix. The subject-static model (C) added subject identity and prefix summary statistics without trial-level dynamics. The individual-history model (D) added subject-specific lag, exponentially weighted, running, and gap features computed prefix-only. The full model (E) additionally included online changepoint features. Three temporal-order controls tested the necessity of chronological structure: time-scrambled (F, within-subject trial-order permutation before feature computation), session-shuffled (G, random session order preserving within-session dynamics), and cross-subject history swap (H, history features from a different subject matched by phase). Paired subject-level comparisons used two-tailed Wilcoxon signed-rank tests (*n* = 17 core subjects), with the paired mean and median AUPRC difference as the effect size and bootstrap 95% confidence intervals (10,000 resamples, seed = 42).

### Subjects

Male Sprague-Dawley rats were purchased from Beijing Huafukang Biotechnology Co., Ltd., Beijing, China, at approximately 6 weeks of age. A total of 25 rats were handled: 24 planned subject positions plus one replacement for the RED 4 position. The animals were acquired in two temporally distinct batches with different identifier schemes: the earlier batch used full-word identifiers (RED/BLACK/BLUE 1-4), whereas the later batch used single-letter identifiers (R/G/B 1-4); RED 4 and R 4 are therefore different subject positions. Primary/core validated behavioral analyses used the 17 subjects that completed all four experimental phases (with both raw event logs and derived trial-level data for parser validation). The frozen reconstructed dataset comprises 24 dataset-level subject labels: 17 core-validated trajectories, 6 raw-only (non-core) trajectories, and the separate RED 4 label; 23 of these trajectories are modelled. Raw-only analyses used the 6 evaluable raw-only subjects; RED 4 is not modelled because none of its 668 phase-coded late-discrimination rows carries an action label (0 of 668; the prediction target is therefore undefined for this dataset-level trajectory). All 6 raw-only animals had experienced periods of incorrect training programming and had initially been set aside from the derived-table four-phase cohort; this group-level history was confirmed by the experimental team, and the surviving records provide illustrative subject-specific details for three. Their raw event logs were retained even though the derived trial-level data required for core validation were unavailable, and all 6 were included in the all-23 robustness analysis rather than excluded from the study or final modelling. The RED 4 label is not one of these six and spans two physical animals (an original animal and a re-trained substitute). Both died before contributing any labelled formal-experiment trials; retained training and phase-coded records include 668 late-discrimination rows, but none carries an action label. This is why 25 animals were handled for 24 frozen dataset-level labels. A subject-identity-cleaned derivative separates the original and substitute as RED 4 and RED 4-sub. This core/extended distinction reflects data-source availability and reconstruction status rather than a post hoc behavioral exclusion criterion. All 24 dataset-level trajectories are reconstructed from the primary Graphic-State operant-chamber logs; the derived per-trial tables produced during the original study serve only as an independent cross-check and exist for the 17 core subjects, where the two records agree (parser validation). Because a data-integrity audit found systematic padding and contamination in those derived tables, they are not treated as ground truth: “raw-only” therefore denotes the absence of a second, independent record to cross-check against, not lower-quality data, and the 6 evaluable raw-only subjects were reconstructed with the same validated parsers. Reconstruction-verification records for these subjects are provided openly in Supplementary Data 9: a per-session raw-file inventory with event-marker counts, a behavioral-plausibility audit (in which every validated decline was rewarded, per the task rule), and the result of rebuilding them from the raw logs with the released parsers; the raw logs themselves are available under controlled access (Data Availability). Rats were socially group-housed three per cage, randomly assigned to cages on arrival, in home cages measuring 460 × 300 × 160 mm each fitted with two 500 mL water bottles. Housing followed a fixed three-per-cage rule: the 17 core validated subjects were housed across six cages and the remaining subjects in separate cages. Where a cage held fewer than three animals, this reflected the two deaths recorded under the RED 4 label before either animal contributed labelled formal-experiment trials, not the raw-only analytical designation or any departure from the housing protocol. Rats were acclimated with 7 days of handling before experiments; cotton nesting/play material was provided as environmental enrichment, and experimenters and caretakers handled and interacted with the animals on an almost-daily basis during the pre-experimental period. This handling-and-enrichment husbandry regime follows the human–rat interaction approach that our group has shown benefits behavioral development and welfare in Sprague-Dawley rats [30]. Environmental conditions were maintained at 20–24°C with a 12h/12h light/dark cycle (lights on at 20:00, off at 08:00; a reversed cycle so that testing occurred during the animals’ dark/active phase) and ad libitum access to water. The animals were fed Beijing Huafukang laboratory rodent chow, with body weight maintained above 90% of free-feeding weight (food was reduced to approximately 80% of ad libitum intake on the day before sessions). Bedding was changed weekly. The reward used in the task was Dustless Precision Pellets Rodent (45 mg, Bio-Serv, #F0021 / #Purified F0021). Among the 17 core subjects at testing, 3 were approximately 20 months old, 2 were approximately 18 months old, and 12 were approximately 6 months old.

### Apparatus

The experiment used six identical sound-insulated Coulbourn Instruments modular shuttle boxes (52.5 × 26.5 × 35 cm each; Holliston, MA, USA), each configured as a wide, single-compartment operant chamber by removing the central liftable gate. The chambers were controlled by Coulbourn Instruments Graphic State software (versions GS3 and GS4). Each chamber contained two retractable response levers flanking a programmable pellet dispenser and infrared-sensed food trough, together with house lights, water access, and a contact (touch) sensor beneath cue lights on the opposite wall. The levers served as the tone-duration classification response options, whereas the contact sensor served as the rewarded decline/give-up option. Figure 1a diagrams the functional arrangement; Supplementary Figure S11 provides a labelled photograph of the original apparatus and preserves its wide shuttle-box configuration.

### Behavioral task, pretraining, and procedure

#### Task overview

Subjects performed an auditory duration-discrimination task in which a pure-tone stimulus had to be classified as “short” or “long” by pressing one of two retractable levers positioned on either side of a central food trough. A contact (touch) sensor mounted on the opposite wall beneath a tri-color cue light provided a rewarded decline/give-up option. Short and long tones were mapped to the two levers, with the mapping counterbalanced across subjects and fixed from pretraining onward (Figure 1a; Supplementary Figure S11).

#### Pretraining

Before formal testing, every subject completed a five-stage shaping sequence: (1) chamber and magazine habituation, with pellets delivered automatically at the food trough; (2) lever-press training, learning to press an extended lever for food; (3) tone-discrimination training, learning to press the lever corresponding to the short versus long tone, with correct presses rewarded and incorrect presses penalized by a delay; (4) contact-sensor training, learning to touch the contact sensor for food when the tri-color cue light was illuminated; and (5) combined-choice training, in which the levers and the contact sensor were simultaneously available after the tone. Each subject accrued at least 50 h of cumulative training before formal testing.

#### Trial structure and timing

Each session began with a 1-min house-light acclimation period. On each trial a pure-tone stimulus (duration 2–8 s; see below) was presented, and at tone offset the trial type was determined and the corresponding manipulanda became available. In forced-lever trials, both levers extended 2 s after tone offset and the contact-sensor cue light remained off (sensor touches were ineffective). In forced-sensor trials, the levers did not extend and the contact-sensor cue light illuminated immediately at tone offset. In choice trials, the contact-sensor cue light illuminated at tone offset and both levers extended 2 s later, so that the animal could either wait and classify the tone by pressing a lever or decline at any time by touching the contact sensor. A correct lever press or a decline (contact-sensor) response was rewarded with food pellets (number set by the reward ratio; see below); an incorrect lever press incurred an additional 45-s wait penalty before the inter-trial interval; and no response within 45 s of tone offset was scored as an omission with no reward. Each trial was followed by a 42-s inter-trial interval. Sessions lasted 120 min, and each subject completed one session per day. In choice-containing test sessions, trial types occurred with probabilities of 25% forced-lever, 25% forced-sensor, and 50% choice (Figure 1b).

#### Stimulus set and difficulty

In the two-stimulus phases (Experiments 1 and 3) the short and long tones were 2 s and 8 s. In the eight-stimulus phases (Experiments 2 and 4) eight equiprobable tone durations were grouped into short (2, 2.44, 2.97, 3.62 s) and long (4.42, 5.38, 6.56, 8 s) categories, defining four difficulty levels of increasing proximity to the category boundary (level 1: 2/8 s; level 2: 2.44/6.56 s; level 3: 2.97/5.38 s; level 4: 3.62/4.42 s).

#### Experimental phases

The behavioral protocol comprised four phases, each run for seven daily sessions per subject and sharing the same 17-subject cohort: Experiment 1 established the choice paradigm with 2s/8s stimuli at a 1:1 lever:sensor reward ratio (one pellet for a correct lever press or a decline); Experiment 2 introduced the eight-stimulus difficulty manipulation at a 1:1 ratio; Experiment 3 manipulated the reward ratio between correct engagement and decline, with a low-decline group (*n* = 9) shifted from 1:1 to 1:2 (favoring decline) and a high-decline group (*n* = 8) shifted from 1:1 to 2:1 (favoring engagement); and Experiment 4 combined the eight-stimulus difficulty manipulation with the between-subject reward-ratio manipulation (1:2, *n* = 9; 2:1, *n* = 8). Because reward-ratio blocks were assigned according to each subject’s earlier decline tendency, reward ratio functioned simultaneously as a current-trial contingency and as part of each animal’s individualized longitudinal history, a central motivation for the triggering-versus-structuring-cause analysis (Figure 1c).

#### Reward contingencies (empirical audit)

An empirical reward-rule audit of the reconstructed data confirmed that validated true decline/give-up actions in discrimination contexts were rewarded in all cases (30,768/30,768; 31 edge-case training/parser rows were excluded from this contingency estimate; Supplementary Figure S4). Incorrect engagements and omissions yielded no reward and triggered a timeout.

#### Data acquisition and preprocessing

Behavior and event timing were recorded automatically by the Graphic State software controlling each chamber. Following the original protocol, a fixed 2,000 ms was subtracted from all lever-press reaction times to correct for the programmed lever-extension delay, and trials exceeding the 45,000 ms program cutoff or lying beyond 2.5 SD of a subject’s mean were removed (<3% of trials). The complete chronological record, spanning all shaping and experimental phases, was reconstructed from the raw Graphic-State logs into the trial-level dataset analyzed here (see Dataset Reconstruction).

#### Dataset reconstruction

The reconstructed dataset comprises 213,990 trial events across 2,162 sessions (spanning up to 117 sessions per subject). Here “2,162 sessions” denotes the number of records in the session-summary table; because the trial-event table’s session_id field indexes raw session fragments, it enumerates more identifiers (2,506), and the dataset-audit usability table lists 2,095 sessions after excluding 67 unusable sessions (session-count reconciliation in Supplementary Data 7). Extensive integrity auditing recovered 9,844 trials across 74 patched sessions (Supplementary Figures S1 and S2; Supplementary Data 2). We restricted training and testing to the 162,304 late-discrimination trials to evaluate forecasting of putative metacognitive give-up behavior on mature behavior (Figure 1d).

#### Trial-flow accounting (CONSORT-style)

- 213,990 trial/event rows (24 subject IDs, 23 modelled, 2,162 sessions) - → 162,304 late-discrimination trials (all subjects) - → 120,300 rows in the model-input/evaluation index; the remaining 42,004 late-discrimination rows fell outside that index - → 48,903 training + 15,320 validation + 51,090 test + 4,987 excluded. Training and validation drew on all 23 evaluable subjects (the 17 core-validated and 6 raw-only subjects, i.e., every subject except RED 4, whose 668 late-discrimination trials carry no action labels); the 51,090 held-out test trials came from the 17 core-validated subjects (test set restricted to core_17).

See Supplementary Information S3 and Supplementary Figure S3 for the full split specification and leakage audit.

### Model architectures and forward-chaining

We utilized a strict chronological 60/40 session split per subject: the first 60% of sessions for each subject comprised the training set, and the final 40% formed the test set. This yielded 51,090 test trials. Models generated step-ahead predictions based strictly on causal prefix history. Because these data are non-exchangeable per-subject streams, we treated leakage-safe chronological evaluation as a substantive methodological commitment rather than a technical detail, given evidence that data leakage pervasively inflates reported performance across machine-learning-based science [31]. We evaluated four major model families: 1. **Static baselines (M1)**: Stimulus-aware logistic regression. Implementations used standard Python machine-learning tooling [32], with hyperparameter screening guided by random-search principles [33]. 2. **Sequential history (M5)**: Gradient-boosted trees (LightGBM) [19] built on lagged actions and EWM averages. 3. **Hidden Markov models (HMM)**: A categorical *K* = 4 HMM [22] fit exclusively on the training prefix, performing online forward-filtering over test observations. 4. **Recurrent neural networks**: GRU [21] (*h* = 4 and *h* = 8 dimensions) equipped with sequence processing and subject embeddings.

### Preregistration

No part of this study was preregistered. The aim is not to confirm or reject a preset hypothesis but to perform history-structured, full-throughput forecasting of outcome behavior (as reflected in the title); the design and analysis are therefore exploratory. Consistent with this, no trials were excluded on the basis of behavioral outcome (correct, incorrect, decline, and omission trials were all retained in the modeling), and the only records set aside were those removed during data-integrity reconstruction (contaminated or padded rows; see Dataset reconstruction). Analyses were scoped to the late-discrimination task phase as described above.

### Use of artificial intelligence and AI-assisted technologies

The central idea of history-structured forecasting and its application to these animal-behavior records were conceived by Bin Yin. The experimental paradigm, animal training and testing, primary data collection, and contemporaneous record-keeping were designed or conducted by the human research team without AI assistance and predated the AI-assisted computational work. A guiding principle was to preserve, reconstruct, and account for every recoverable individual record rather than discard procedurally irregular or inconvenient trajectories. Six animals that had experienced periods of incorrect training programming were initially set aside from the 17-subject core cohort but were subsequently included in the all-23 robustness analysis. RED 4 was separate from these six: the dataset label covered an original animal and its replacement, neither of which contributed a labelled formal-experiment trial, and its retained phase-coded records lacked the action labels required to define the forecasting target. After data collection, OpenAI GPT-5-series systems (including Codex), Anthropic Claude Code, and Edison Scientific Kosmos were used under author direction to assist with reconciling legacy data structures; implementing and executing author-specified cleaning, forecasting, and validation code; conducting reproducibility and consistency checks; organizing the release packages; and refining language. Their outputs were treated as unverified until checked against source records, versioned data, executable code, independent reruns, and cross-tool comparisons. Quantitative figures and tables were rendered from validated data through transparent, version-controlled scripts in documented programming environments; they were not directly generated by AI systems. The apparatus image is an original experimental photograph, and Figure 1 is a scripted schematic grounded in the documented hardware and procedures; neither is synthetic imagery. AI tools did not autonomously determine the hypotheses, final model acceptance, interpretation, or conclusions. The human authors determined the research questions, final model specifications, analytical decisions, evidential thresholds, interpretations, and final wording. Unsupported or non-reproducible results were corrected, withdrawn, relabelled as archival or exploratory, or reported as limitations. No AI system was treated as an author, and the human authors reviewed and approved all outputs and take full responsibility for the work.

## Supporting information

Supplementary Data ZIP

## Ethics Statement

All experimental procedures involving animals were approved by the Experimental Animal Ethics Committee of Fujian Normal University (protocol IACUC-20230055) and were conducted in accordance with institutional guidelines for animal care and use and with the Chinese national standard GB/T 35892-2018 (“Laboratory animal—Guideline for ethical review of animal welfare”). At the end of the study, animals were euthanized by carbon dioxide (CO₂) inhalation.

## Acknowledgements

We thank the experimental animals whose behavior made this study possible. We also thank the members of the laboratory who assisted with animal care, behavioral testing, and data organization.

## Author Contributions

- **Bin Yin**: Conceptualization, Methodology, Resources, Software, Formal analysis, Data curation, Writing – original draft, Writing – review & editing, Visualization, Supervision, Project administration
- **Ya-Xin Wang**: Investigation, Methodology, Data curation, Writing – review & editing
- **Chongyi Liu**: Data curation, Validation
- **Liya Fu**: Data curation, Validation

## Data Availability

The de-identified trial-level dataset and model-output tables are available in Zenodo version v1.18 at https://doi.org/10.5281/zenodo.21882292 (all versions: https://doi.org/10.5281/zenodo.21228501) under a CC BY 4.0 license [34].

Supplementary Information provides supporting methods and analyses.

The Supplementary Data PDF provides a reader-facing index and rendering of Supplementary Data 1-9.

The Supplementary Data ZIP contains the corresponding machine-readable files and descriptions.

Reconstruction and verification records for the raw-only subjects (raw-file inventory, behavioral-plausibility audit, and raw-to-trial reconstruction results) are openly available as Supplementary Data 9 and within the same Zenodo version (audit_documentation/). The raw Graphic-State event logs and lab-specific metadata are available under controlled access from the corresponding author, during peer review and after publication, for the purpose of reproducing or extending the reported analyses; access is restricted because these files contain operational annotations not suitable for unrestricted redistribution. Requests will receive a response within two weeks and, if approved, require a data-use agreement specifying permitted analyses, confidentiality obligations, and no redistribution of the raw files.

## Code Availability

All analysis code is available in the same Zenodo v1.18 record under a BSD-3-Clause license [34]. The repository reproduces the M5+Changepoint teacher-forced benchmark and the eight-model structuring-cause validation, including the per-subject Wilcoxon tests and bootstrap confidence intervals behind Figure 2 and Supplementary Data 5. On the pinned stack (LightGBM 4.6.0) the pipeline is deterministic and the values reported here are exactly what it produces: the reported benchmark (action accuracy 51.7%, log loss 1.098, decline AUPRC 0.525, temperature 1.347 on 51,090 held-out trials) and the structuring-cause contrasts are the released values, verified by an included leakage audit and a fail-closed verification script (verify_reproduction.py). The earlier frozen model-output tables are retained only as a historical record. The action-conditioned reaction-time analysis (06_rt_model.py) and the c_t structuring-state profiles (08_ct_profiles.py) are provided as released code; because the original reaction-time derivation used a different subject population and the c_t profiles use simplified definitions, their archival values are retained for those specific tables and are labelled as such (release manifest, Supplementary Data 4). The closed-loop simulation (07_free_running_simulation.py) is an exploratory analysis that reproduces the archived over-decline direction and magnitude (+17.1 pp, 0/115 collapses) on retraining, and is not primary evidence. A separate, controlled-access package documents the raw-data reconstruction: the standard-session reconstruction is executable, while the patched sessions and the finalized dataset are provided as frozen, validated outputs.

