## Supplementary Information for "History-structured forecasting of rewarded give-up behavior in a rodent metacognition task"

This supplement supports the manuscript "History-structured forecasting of rewarded give-up behavior in a rodent metacognition task." It emphasizes auditability, leakage control, negative results, and complete per-subject reporting.

#### S1. Dataset reconstruction, parser validation, and anomaly audit

##### S1.1 Raw data provenance

- Raw event log files: 3,066.
- Frozen canonical trial/event dataset: 213,990 rows.
- Canonical session summary: 2,162 sessions total (1,938 scheduled + 224 unscheduled).
- Subjects: 24 subject IDs (17 core-validated + 6 raw-only + RED 4); 23 modelled (RED 4 has no action labels); 25 animals handled.
- Late discrimination subset used for primary modeling: 162,304 trials.

**Session-count reconciliation note:** Supplementary Data 1 summarizes the 2,095 sessions retained in the dataset-audit usability table. The full canonical session summary contains 2,162 sessions. The 67-session difference corresponds exactly to sessions flagged modeling_use =

EXCLUDE in the canonical session summary. These excluded sessions were scheduled records retained in the canonical trajectory for provenance completeness but were not included in the usability/audit table because they were review-needed or excluded sessions, most commonly no-raw-file scheduled entries, no-experiment rows, auxiliary protocol records, and known wrong-program/home-cage irregularities. The complete session-by-session reconciliation is provided in Supplementary Data 7.

##### S1.2 Parser validation

Parser validation achieved 97.9% effective accuracy across 583 sessions (exact plus terminal-edge matches, 571 of 583; Supplementary Figure S2); counting the 6 additional multi-session sessions as usable yields the 99.0% reported in Supplementary Data 1's `effective_accuracy_pct` column. Supplementary Data 1 documents the number and percentage of sessions passing exact or effective parser checks, known edge cases, and interpretation.

##### S1.3 Patched sessions

74 sessions required patching, recovering 9,844 trial rows (documented in Supplementary Data 2). These patched sessions were retained only after rigorous integrity review.

##### S1.4 Missingness and anomaly handling

Extreme RTs (>60 s) and zero RTs were retained with explicit flags. No RT clipping, winsorizing, or imputation was applied. Sessions with unresolved protocol/counterbalance conflicts were excluded from accuracy-based analyses when mapping correctness could not be trusted. Supplementary Figure S1 shows the trial recovery and filtering flow.

### S2. Task mapping, counterbalance, and reward-rule audit

#### S2.1 Sensor-to-choice mapping

7 sessions (0.2%) were flagged with unresolved protocol versus counterbalancing conflicts and excluded from accuracy-based analyses (reconstruction QC report and response-mapping audit in the Zenodo deposit `audit_documentation/`).

#### S2.2 Reward-rule audit

Authoritative rules: 1. **Decline/Give-up = rewarded**. True decline actions enter a reward/feed route. 2. **Correct engage = rewarded**. 3. **Incorrect engage = not rewarded; timeout**. 4. **Omission = not rewarded; timeout**.

Empirical confirmation (Supplementary Figure S4): 30,768/30,768 validated true decline/give-up actions were rewarded (100%); 31 apparent exceptions were parser/training-context artifacts and were excluded from this count.

Reward magnitude: - Standard discrimination contexts: decline = 1 pellet, correct engage = 1 pellet. - 1VS6 context: decline = 1 pellet, correct engage = 6 pellets.

#### S2.3 Reward-ratio block design

The experimental design included reward-ratio blocks that modulated the relative value of lever-based correct engagement and contact-sensor decline. The three main reward-ratio contexts were:

- **1:1 baseline**: Correct lever engagement and decline/contact-sensor each yielded one pellet when rewarded.
- **High-engagement reward (e.g., 2:1)**: Assigned to high-decline animals. Correct lever engagement yielded increased reward (e.g., 2 pellets) relative to decline (1 pellet), incentivizing engagement over the give-up option.
- **High-decline reward (e.g., 1:2)**: Assigned to low-decline animals. Decline/contact-sensor yielded increased reward (e.g., 2 pellets) relative to correct lever engagement (1 pellet), increasing the relative value of the give-up option.

Mixed contexts combined reward-ratio manipulations with stimulus-difficulty manipulations (e.g., 1VS6 contexts where correct engagement yielded 6 pellets).

Reward-ratio blocks were assigned based on prior individual decline tendency: animals with higher decline rates received the high-engagement reward context, and animals with lower decline rates received the high-decline reward context. This assignment procedure means that the reward-ratio condition was both a current triggering context and a historically contingent environmental manipulation reflecting each subject's prior behavioral trajectory.

##### S2.4 Role of the reward pathway in the simulation

The over-decline drift depends on the rewarded decline pathway. When simulated decline actions are rewarded, as they are in the task, the reward-history feedback loop strengthens and the over-decline tendency grows; when decline actions are not rewarded, the drift is muted. Within this archived simulation, the reward rule modulates the modeled drift; this diagnostic does not show that reward-history features, rather than recent action history, drive the primary teacher-forced forecast. The exact magnitude is exploratory and configuration-sensitive (see S7) and is not used as primary evidence.

### S3. Forecasting protocol, chronological splits, and leakage audits

#### S3.1 Primary split

- Split type: strict chronological 60/40 session split per subject.
- Training data: first 60% of sessions per subject.
- Test data: final 40% of sessions per subject (Supplementary Figure S3).
- Primary held-out test set: 51,090 trials.
- Primary outcome: 4-action prediction (left, right, decline, omit).
- Chance level: 25%.

#### S3.2 Operationalizing triggering versus structuring causes

Following the triggering/structuring-cause framework (Dretske [17]; Potter and Mitchell [18]), feature inputs are grouped into two theoretical classes (Supplementary Figure S9): - **Triggering causes** (current-trial features): stimulus duration, reward-ratio context, context flags, rule contingencies. Used exclusively in the M1 static baseline. - **Structuring causes** (prefix-derived features): lagged actions, exponentially weighted moving averages, cumulative reward histories, inter-session gap durations, changepoint segment age, rolling delta statistics, and $c_t$ profile features. These represent historically accumulated constraints.

##### S3.3 Leakage-audit checklist

All history features were computed from causal prefixes only. A nine-point leakage audit verified: 1. Chronological split performed by session order within subject. 2. Feature engineering uses prefix-only histories. 3. Rolling/EWM windows shift by one trial where required. 4. Test labels excluded from model training. 5. Calibration temperature fit on a pre-test validation partition only. 6. HMM fitted on train prefix only. 7. HMM test predictions generated by online filtering, not train+test Viterbi paths. 8. Free-running simulation histories use simulated prior actions/rewards only after start. 9. Per-subject summaries computed after prediction and not used as predictors.

#### S4. Calibration and full model-comparison details

##### S4.1 Primary model (M5+Changepoint)

Temperature-scaled LightGBM: - Features: 64. - Test trials: 51,090. - Action accuracy: 51.7%. - Log loss: 1.098. - Macro-F1: 0.476. - Decline AUPRC: 0.525 (a lift over the 0.30 decline prevalence in this test set). - Decline F1: 0.424. - Decline ECE: 0.036. - Temperature: $T = 1.347$.

##### S4.2 Model comparison and calibration

Supplementary Data 2 provides the full model comparison across all baseline and sequential architectures. Temperature scaling was preferred because it improved multiclass log loss (1.138 to 1.098) while maintaining rank ordering; it slightly worsened decline ECE (0.0327 to 0.0363), so the calibration benefit rests on log loss and Brier score rather than on ECE.

**Evaluation-set size caveat:** Models in Supplementary Data 2 were evaluated on different eligible-trial sets across analysis phases (e.g., GRU models: 39,237 trials; M5+Changepoint: 51,090 trials). The final M5+Changepoint benchmark and the structuring-cause validation use fixed evaluation indices. Where evaluation sets differ, comparisons support model-family screening rather than exact paired superiority claims.

#### S5. HMM leakage correction and proper forward-chained evaluation

An initial HMM analysis reported decline AUPRC 0.629. This value was inflated because Viterbi decoding used train+test sequences, allowing test-set information to influence inferred state paths. Proper train-only categorical $K = 4$ HMM evaluation yielded an action accuracy of 48.6% and decline AUPRC of 0.440, falling well below the transparent M5 floor (Supplementary Figure S5). The HMM's near-absorbing disengaged state (mean dwell 928 trials) helps explain its poor forward prediction.

HMM states are descriptive empirical clusters of sequential choice behavior, not psychological or neural states.

#### S6. GRU and recurrent-model negative results

The best recurrent model (GRU $h = 8$ with embeddings and changepoint features) achieved 51.9% action accuracy, a log loss of 1.132 and decline AUPRC of 0.533 on 39,237 eligible trials. The M5+Changepoint model achieved 51.7%, 1.098 and 0.525, respectively, on 51,090 trials.

Because the eligible-trial sets differ, these values support model-family screening rather than an exact paired superiority claim.

Furthermore, the archival GRU reaction-time result had an MAE of approximately 7.4 s, compared with 1.43 s standalone and 1.26 s integrated for the dedicated action-conditioned RT model (Supplementary Figure S6). Taken together with the smaller GRU evaluation set, these results motivated retention of the transparent history model; they are not an exact matched-set superiority test.

#### S7. Exploratory closed-loop simulation: mechanism and robustness

The free-running closed-loop simulation feeds the model's own predictions back as ongoing history, probing the boundary between step-ahead forecasting and behavioral generation. We report it as an exploratory, configuration-sensitive analysis rather than as primary evidence.

**Mechanism within the archived simulator.** In the recovered Phase-3F implementation, rewarded predicted declines update rolling reward-history features and can amplify later simulated decline probabilities. Holding those rolling features fixed attenuated the archived drift. This identifies a feedback pathway within that exploratory simulator; it does not establish that reward-history features are the load-bearing signal in the primary teacher-forced forecast, for which the current ablation instead identifies recent action history as critical.

**Result (archived Phase-3F outputs).** The archived Phase-3F simulation tables, retained as the historical record for this exploratory analysis, show a source-dependent over-decline of approximately +16.5 to +18.6 pp across seeds, with no collapsed runs. The magnitude depends on the simulation configuration (feature set, number of seeds, and feature-update rule), so these archived tables, not any single re-run, are the reference, and Supplementary Figure S12 is drawn from them. Full provenance of the archived outputs is documented in the data package's audit documentation.

**Per-subject reliability.** Free-running generation reliability varies across subjects: of the 23 evaluable subjects, 2 are high-, 4 medium-, and 17 low-reliability, so most subjects show a substantial gap between teacher-forced prediction and closed-loop generation. This reinforces treating the closed-loop analysis as exploratory rather than as a per-subject synthetic generator.

**Conclusion.** The closed-loop simulation is exploratory and configuration-sensitive. The primary reproducible results are the teacher-forced M5+Changepoint benchmark and the structuring-cause validation. See Supplementary Figure S12.

### S8. Per-subject forecast summaries and individual profiles

#### S8.1 $c_t$ profile table

Five non-tautological $c_t$ features (Supplementary Data 3) demonstrated distinct individual structuring strategies: - transition_rate: 0.565. - context_adaptation_speed: 0.146. - win_stay_rate: 0.574. - lose_shift_rate: 0.382. - gap_sensitivity: 0.184.

Win-stay rate showed a suggestive positive association with simulation decline bias ($r$ = +0.483, $p$ = 0.019, $n$ = 23; not FDR-corrected).

#### S9. Structuring-cause validation

To test whether individual diachronic history carries structuring-cause-like predictive information for trial-by-trial behavior, we trained eight LightGBM multiclass models spanning a 2×2 taxonomy (group-level vs individual-level × synchronic vs diachronic) plus three temporal-order controls (time-scrambled, session-shuffled, cross-subject swap).

Absolute accuracies in this validation analysis (Supplementary Data 5) exceed the primary M5+Changepoint accuracy (51.7%) because this analysis included additional triggering-feature covariates (trial type, paradigm, context); all conclusions are based on within-analysis contrasts and paired subject-level comparisons (two-tailed Wilcoxon signed-rank tests, $n = 17$ core subjects, 10,000 bootstrap resamples). The two primary decline-AUPRC contrasts, D-vs-A and D-vs-B, each had an exact $p = 1.53 \times 10^{-5}$ and remain below .001 after conservative correction across the eight reported model contrasts. Per-subject deltas, Wilcoxon statistics, exact $p$-values and bootstrap 95% confidence intervals for every contrast are tabulated in the statistical_tests section of Supplementary Data 5.

Key Paired Comparisons:

- **Individual-history vs Triggering-only:** Decline AUPRC +0.129 [0.074, 0.186], $p = 1.53 \times 10^{-5}$; all 17 subjects improved (Supplementary Data 5, Supplementary Figure S7).
- **Individual-history vs Group-history:** Decline AUPRC +0.137 [0.081, 0.195], $p = 1.53 \times 10^{-5}$; all 17 subjects improved.
- **Individual-history vs Subject-static:** Decline AUPRC +0.126 [0.070, 0.184], $p = 4.58 \times 10^{-5}$; 16 of 17 subjects improved.

- **M5+Changepoint vs Individual-history:** Decline AUPRC +0.0036 [0.0006, 0.0075], $p$ = .080; the small increment did not reach significance.
- **Depth-matched chronological vs Time-scrambled:** Decline AUPRC +0.118 [0.062, 0.176], $p$ = 7.63 × 10^-5 (Supplementary Figure S8).
- **Depth-matched chronological vs Session-shuffled:** Decline AUPRC +0.008 [0.001, 0.015], $p$ = .071; the small difference was borderline.
- **Depth-matched chronological vs Cross-subject swap:** Decline AUPRC +0.099 [0.052, 0.149], $p$ = .000214.
- **Full-stream vs depth-matched chronological history:** Decline AUPRC +0.0003 [-0.0048, 0.0046], $p$ = .284, supporting the intended history-depth null.

**Choice-only robustness (deterministic-trial control).** Every model in this validation includes the current trial type as a covariate. On forced-sensor trials the contact sensor is the only available response, so a decline is near-mandatory (decline rate 0.77 in the held-out test set), and trial type therefore near-determines the decline label there; other non-choice trial types likewise constrain the available response. This inflates the absolute decline AUPRC of all eight models (e.g. triggering-only Model A reaches 0.684 pooled). To verify that the individual-history advantage is not driven by mandated or otherwise deterministic trial types, we recomputed the D-vs-A decline AUPRC restricted to the 25,036 genuine choice trials (decline base rate 0.213). The advantage was larger, not smaller: pooled Model A 0.208 (essentially the choice-trial prevalence) versus Model D 0.671, a per-subject mean difference of +0.253 (median +0.292; 16 of 17 subjects; exact Wilcoxon $p = 4.6 \times 10^{-5}$), compared with +0.129 on all trials. Because the restriction removes all non-choice categories rather than forced-sensor trials alone, it establishes that deterministic non-choice trials do not manufacture the individual-history signal and collectively dilute its within-choice contrast. Script 11 emits both the summary (choice_only_robustness.csv) and the 17 paired subject-level values (choice_only_per_subject.csv), which the fail-closed verifier reconciles independently.

#### S10. Exploratory unseen-cohort transfer and irregular-history stress test

As an exploratory parameter-transfer probe, we fitted the M5+Changepoint model on the 17 core validated subjects only and applied it to six raw-only subjects (BLACK 1, BLACK 3, BLUE 1, BLUE 3, RED 1, RED 3), using each target animal's own prior behavioural history as input. Because each target contributes its own earlier history, this is a small ($n = 6$) exploratory probe rather than a clean unseen-cohort generalization test. RED 4 was not evaluable because all its late-discrimination trials had missing action labels.

The experimental team confirmed that all six raw-only animals had experienced periods of incorrect training programming and had consequently been set aside from the original core cohort. Surviving records provide illustrative subject-specific details for three: RED 3 was placed in the wrong home cage, resulting in a wrong experimental program during one training stage; BLUE 1 received an extra session in stage 8; and RED 1 had an anomalous session in stage 7. All six remained separate from RED 4 and were included in the all-23 robustness analysis because their histories and action labels were recoverable.

As an exploratory parameter-transfer probe, we fitted the M5+Changepoint model on the 17 core animals only (parameters and temperature from core training/validation) and evaluated it on the six raw-only animals' held-out final-40% suffix (4,987 trials), using each target's own chronological prior history as input (scripts/13). The transfer model reached 58.5% pooled four-action accuracy and beat a last-action persistence baseline for all six animals; however, it exceeded each animal's suffix modal-action rate in only one of six, so nominal 25% chance is not an adequate benchmark here. Treating the subject as the inferential unit ($n$ = 6), the individual-history model did not reliably outperform the triggering-only model (mean accuracy difference 0.050, exact sign-flip $p$ = .94; mean decline-AUPRC difference 0.008, $p$ = .59), and no order effect was detected (time-scrambled versus chronological history, not significant).

Replacing each target's history with a donor animal's sharply reduced action accuracy for all six subjects (exact $p$ = .03), but this control directly perturbs lagged-action predictors and did not significantly change decline AUPRC ($p$ = .44). These findings are descriptive and exploratory, from six animals, and are not evidence of broad cross-subject generalization; the primary robustness result is the within-subject all-23 forecast (Figure 6), in which the model beat every raw-only subject's prefix-derived modal action and last-action persistence (and the conservative held-out modal rate in one of six).

#### S11. Reproducibility package and data/code release manifest

The reproducibility manifest (Supplementary Data 4) enumerates the released canonical dataset and model-output tables. On the pinned LightGBM 4.6.0 stack, the released code deterministically reproduces the teacher-forced M5+Changepoint benchmark (51.7% accuracy, log loss 1.098, decline AUPRC 0.525, $T$ = 1.347; 51,090 test trials) and the eight-model structuring-cause validation, including the per-subject tests and bootstrap intervals in

Supplementary Data 5. A leakage audit and fail-closed verifier are included. Earlier frozen model outputs are retained as historical records; archival values remain explicitly labelled only for the reaction-time analysis, the simplified *c_t* profiles and the exploratory Phase-3F context. The raw-data reconstruction is documented separately with provenance materials and frozen outputs; that package is a provenance record rather than a fully executable generator for every patched raw session.

A subject-identity-cleaned copy of the dataset is additionally provided in the open Zenodo v1.18 data deposit (https://doi.org/10.5281/zenodo.21882292; all versions: https://doi.org/10.5281/zenodo.21228501), reproducible with the released subject-identity-cleaning script. It splits the `RED 4` identifier, which fused a deceased original animal and a substitute run in its place, into `RED 4` and `RED 4-sub` (this separates the identities but does not recover labels: all 668 of these late-discrimination trials remain without an action label, so RED 4 stays outside the forecasting analyses), and removes 99 early-shaping trials that were sourced from another animal's files, so that every subject identifier corresponds to a single physical animal. Primary reported metrics were computed from the modeling inputs; the subject-identity-cleaned v1.6 copy is provided for future identity-sensitive longitudinal analyses and does not alter the primary teacher-forced benchmark or structuring-cause validation, because the affected rows lie outside the late-discrimination modeling index. The original canonical dataset is retained as the primary release for exact reproducibility of the published numbers. A subject-identity-cleaning audit report accompanies the cleaned copy in the data deposit.

#### S12. Session-count reconciliation

Supplementary Data 7 provides the complete session-by-session reconciliation of the 67-session difference between the canonical session summary (2,162 sessions) and the dataset audit usability table (2,095 sessions). The reconciliation directory contains:

- excluded_67_sessions_reconciliation.csv: Full list of 67 excluded sessions with metadata (subject, session ID, date, phase, protocol, schedule status, modeling use flag).
- excluded_67_by_phase.csv: Excluded session counts by training phase.
- excluded_67_by_subject.csv: Excluded session counts by subject.
- excluded_67_reason_summary.csv: Excluded sessions grouped by schedule status and protocol.
- session_count_reconciliation_note.md: Summary note documenting the reconciliation logic.

All 67 sessions were flagged `modeling_use = EXCLUDE` in the canonical session summary. The most common exclusion reasons were: no-experiment entries (16 sessions), 8-stimulus protocol sessions (12), food-competition probes (10), 1VS1 tone test sessions (12), equipment downtime/no-raw-file entries (8), and lever/balance test sessions (4).

#### Supplementary Data

The following data files accompany this article. They are curated tables tied to specific results; the complete dataset, all model outputs, and the reconstruction code are in the open Zenodo v1.18 deposit (https://doi.org/10.5281/zenodo.21882292).

- **Supplementary Data 1**. Dataset audit: per-session usability table for the canonical dataset.
- **Supplementary Data 2**. Model comparison across all baseline, sequential, changepoint and recurrent architectures.
- **Supplementary Data 3**. Individual structuring-state ($c_t$) features per subject.
- **Supplementary Data 4**. Reproducibility manifest for the released canonical dataset and model-output tables.
- **Supplementary Data 5**. Structuring-cause validation: per-subject results for the eight-model taxonomy and temporal-order controls.
- **Supplementary Data 6**. Exploratory unseen-cohort transfer: transfer of the core-trained model to the six raw-only subjects.
- **Supplementary Data 7**. Session-count reconciliation: session-by-session accounting of the 2,162 vs 2,095 difference (compressed folder).
- **Supplementary Data 8**. All-23-subject evaluation: per-subject action accuracy and decline AUPRC under the leakage-safe 23-subject split.
- **Supplementary Data 9**. Raw-only subject reconstruction and verification: raw-file inventory, behavioral-plausibility audit, and raw-to-trial reconstruction results (compressed folder).

These files differ from the Zenodo deposit in scope: they are focused tables supporting specific claims and figures, whereas the Zenodo record is the complete, independently citable archive for full reproduction.

### Supplementary Figures

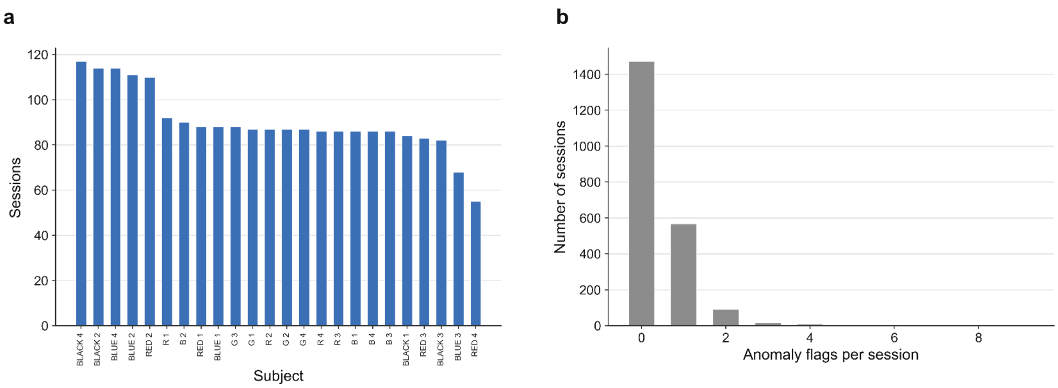


**Figure S1. Sessions per subject and anomaly-flag distribution.** Trial recovery and filtering flow from raw event logs to the final canonical dataset.

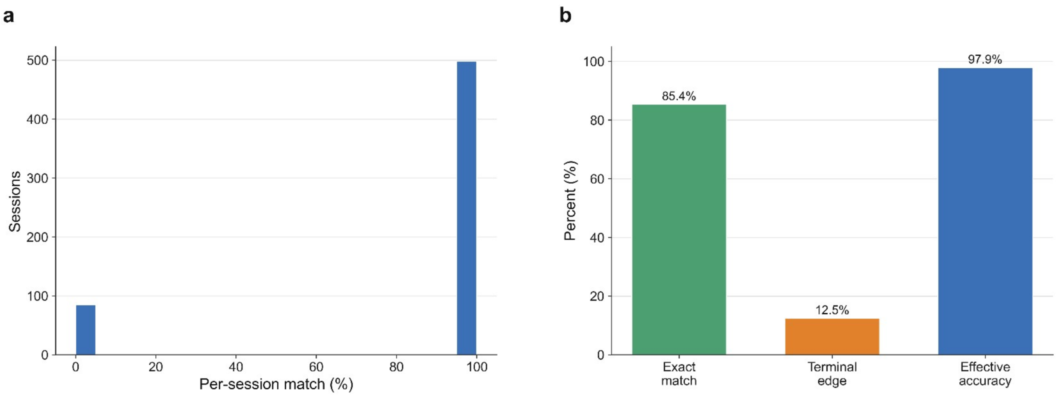


**Figure S2. Parser validation.** 85.4% exact match, 12.5% terminal-edge, 97.9% effective accuracy ($N = 583$ validated sessions).

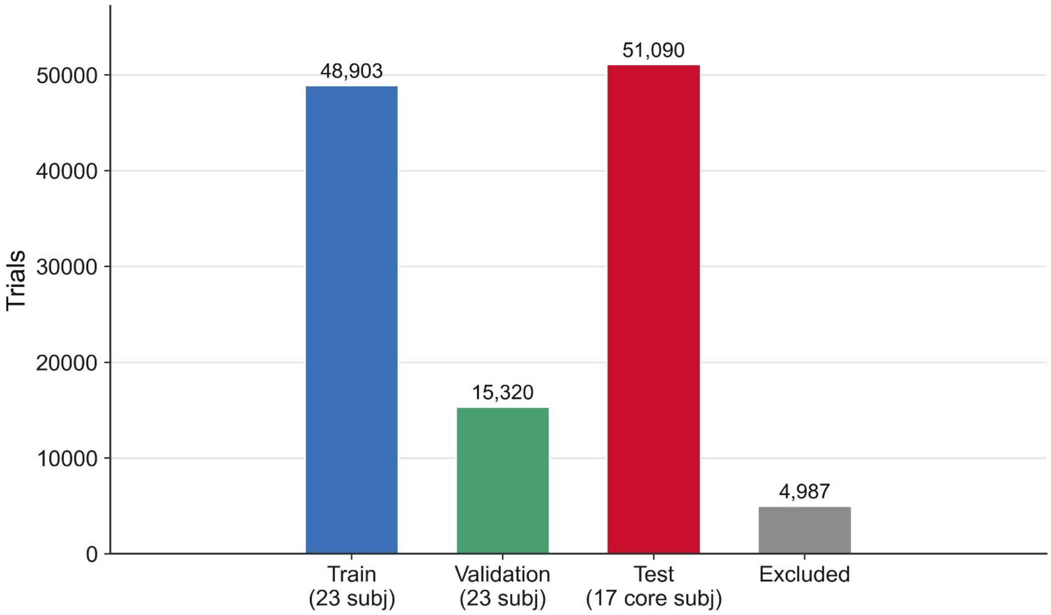


**Figure S3. Chronological prefix-to-suffix split.** 48,903 training + 15,320 validation + 51,090 test trials (120,300 index rows; zero future-leakage verified).

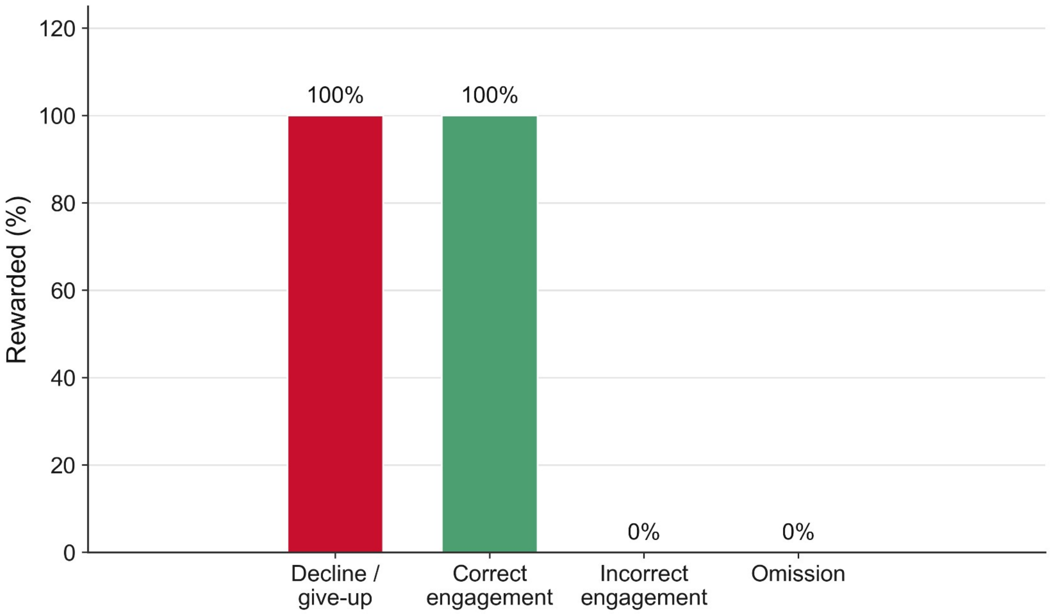


**Figure S4. Reward-rule audit.** 30,768/30,768 validated true decline/give-up actions were rewarded; 31 edge-case training/parser rows were excluded from this count.

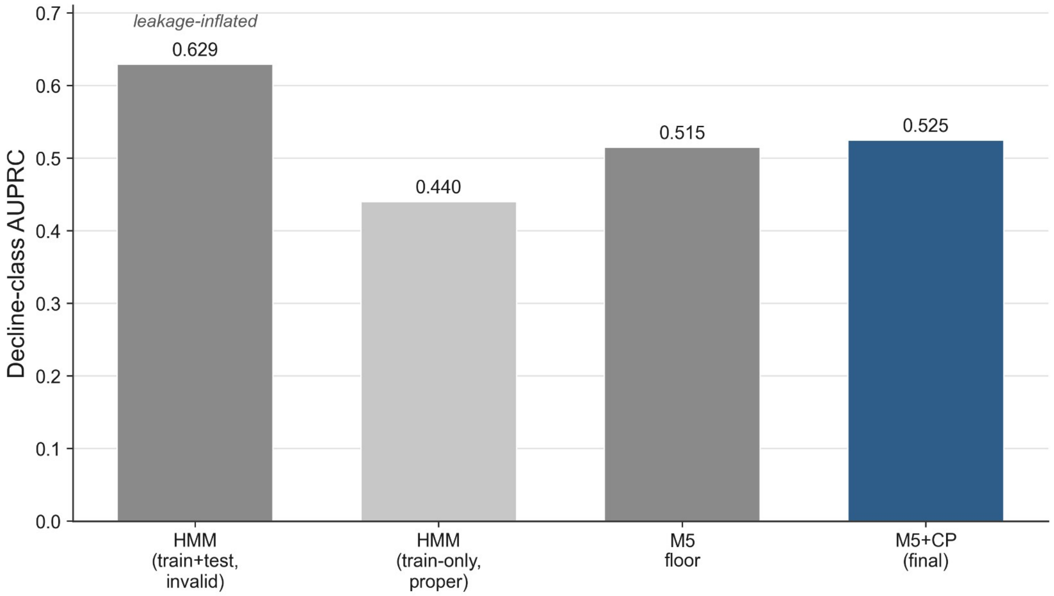


**Figure S5. HMM and changepoint diagnostics.** The 0.629 decline AUPRC obtained from invalid train-plus-test Viterbi decoding was leakage-inflated. Proper train-only evaluation yielded 0.440, below the archival M5 floor (0.515) and the fresh final M5+Changepoint result (0.525).

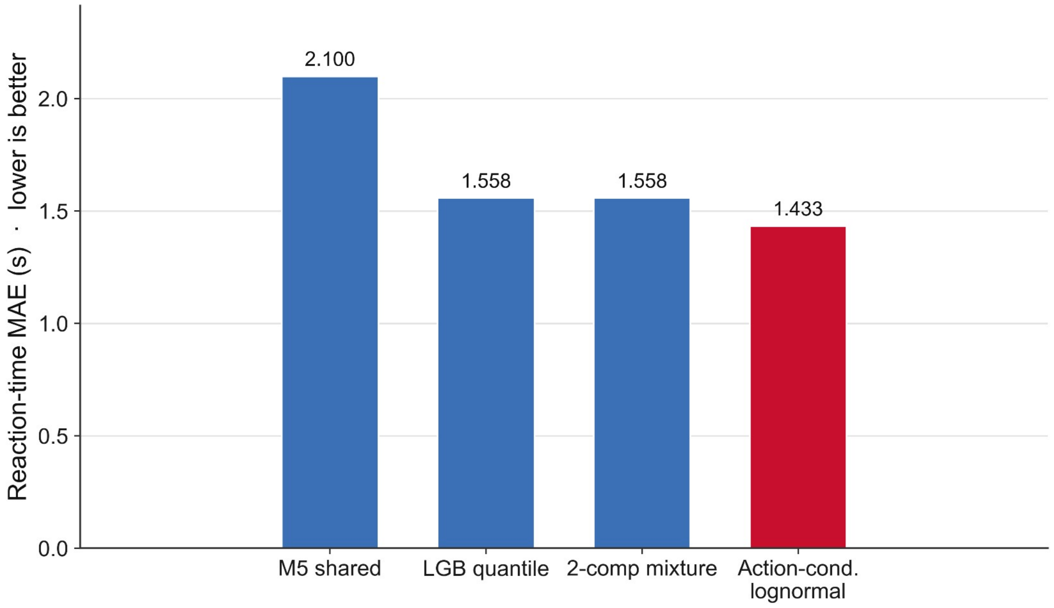


**Figure S6. Dedicated reaction-time model performance.** The action-conditioned lognormal RT model achieved 1.26 s integrated MAE (1.43 s standalone), substantially outperforming the Phase 3B baseline (2.10 s) and the GRU RT head (7.4 s).

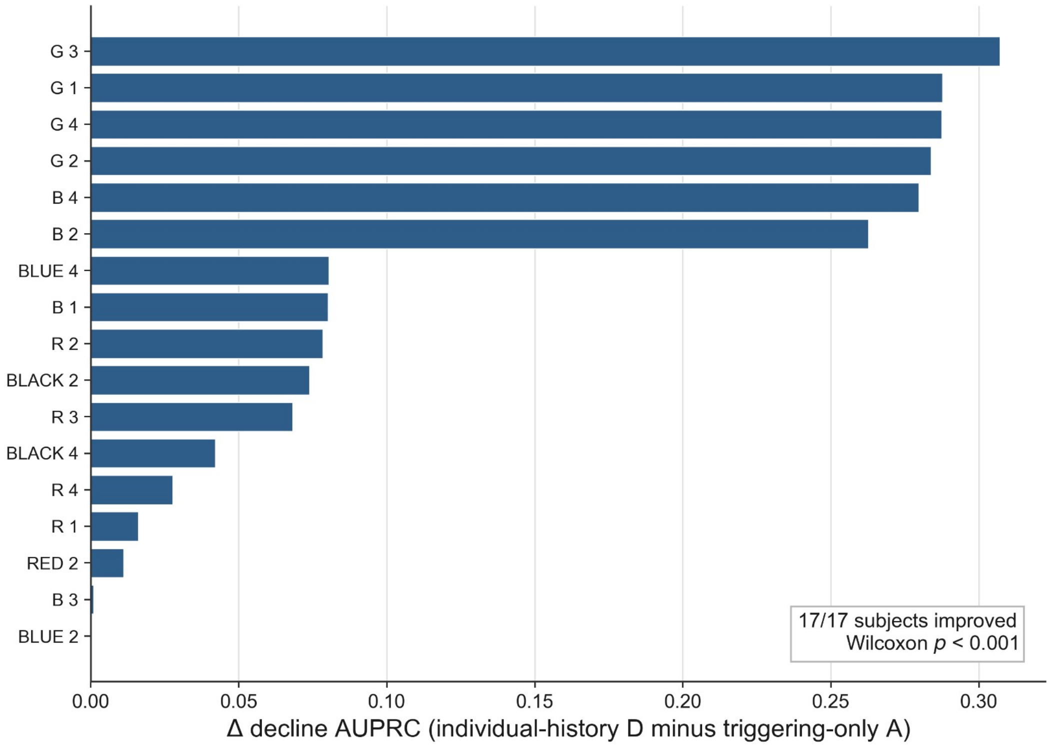


**Figure S7. Per-subject individual-history advantage.** Per-subject decline-AUPRC difference between Model D (individual-history) and Model A (triggering-only), ordered by magnitude. All 17 subjects improved; the paired mean difference was +0.129 (bootstrap 95% CI [0.074, 0.186], Wilcoxon $p < .001$; Supplementary Data 5).

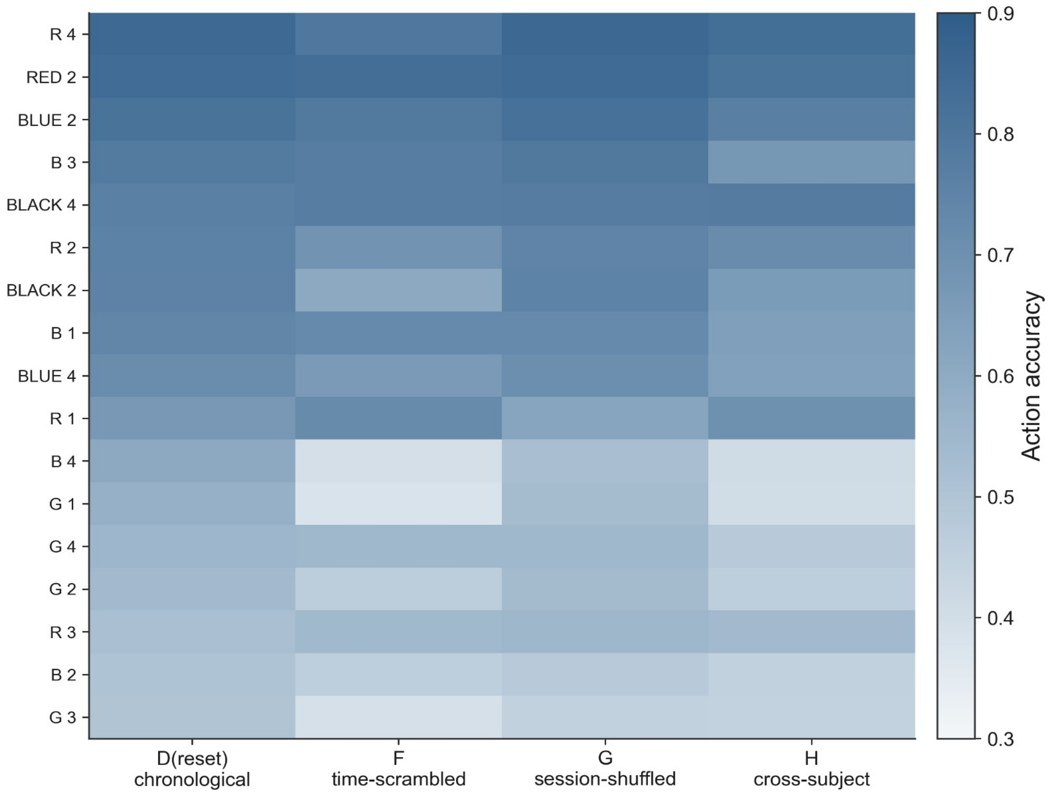


**Figure S8. Diachronic-order control heatmap.** Per-subject action accuracy for the depth-matched chronological anchor D(reset), time-scrambled F, session-shuffled G and cross-subject-swap H. The paired inferential tests use the depth-matched anchor and are reported in Supplementary Data 5.

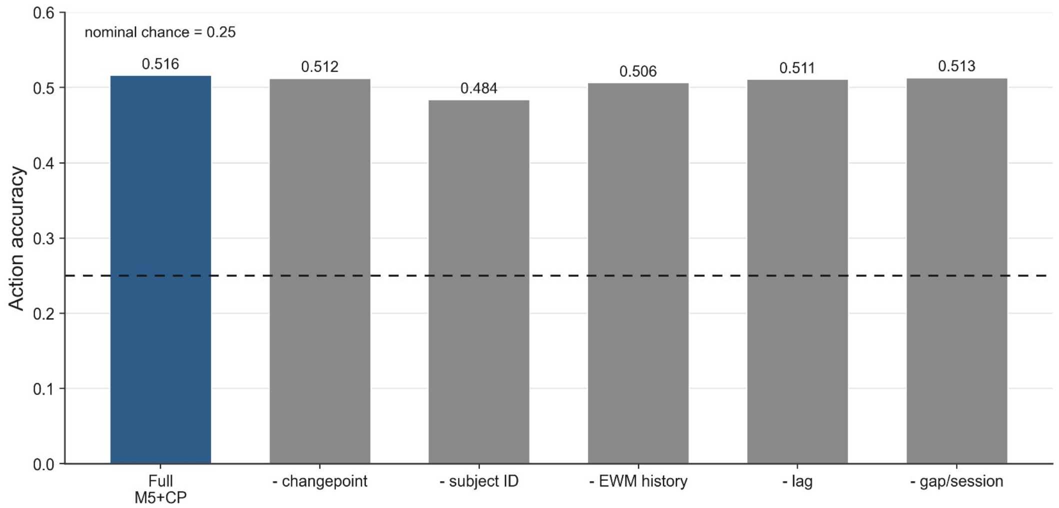


**Figure S9. Archived Phase-3 feature-group ablation.** Action accuracy for the original full M5+Changepoint model and variants omitting one feature group. Removing subject identity produced the largest reduction; omitting changepoint, EWM-history, lag or gap/session features produced smaller reductions. These descriptive archival screening values were not rerun on the canonical stack and are not part of the primary reproducible comparison.

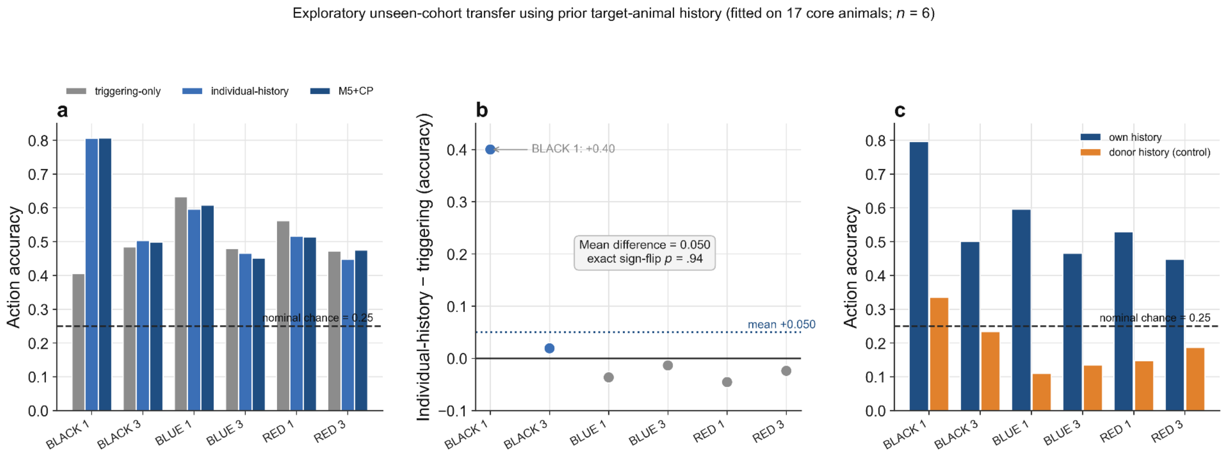


Figure S10. Exploratory unseen-cohort transfer with prior target history. Models fitted to the 17-animal core cohort, evaluated on the chronologically final 40% of trials from the six raw-only animals using each target's own prior behavioural history. (a) Per-subject action accuracy across triggering-only, individual-history and M5+Changepoint. (b) Per-subject individual-history-minus-triggering differences; the mean difference is 0.050 (exact sign-flip $p$ = .94), driven by one animal (BLACK 1, +0.40). (c) Substituting a donor animal's history collapses accuracy, a control that directly perturbs lagged-action predictors. Exploratory and underpowered ($n$ = 6); Supplementary Data 6.

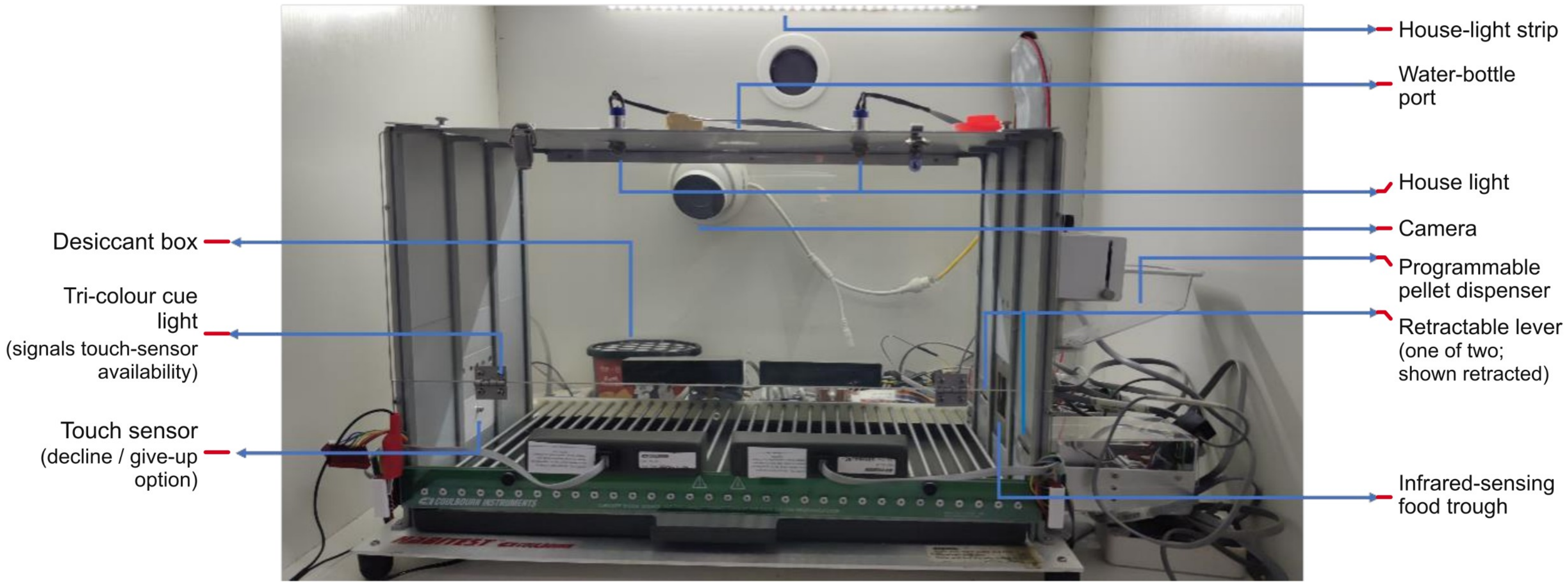


**Figure S11. Photograph of the customized shuttle-box apparatus.** Photograph of one of the six identical Coulbourn Instruments modular shuttle boxes (52.5 × 26.5 × 35 cm) configured as a wide, single-compartment operant chamber by removing the central liftable gate (thesis Fig. 3-1). Labelled components include the camera, desiccant box, tri-colour cue light (signals contact-sensor availability), contact sensor (the rewarded decline/give-up option), house-light strip, water-bottle port, house light, programmable pellet magazine/hopper, response lever (shown retracted) and infrared-sensed food trough. The apparatus contained two retractable response levers, one on each side of the trough. Short/long tone-to-lever mapping was counterbalanced across subjects.

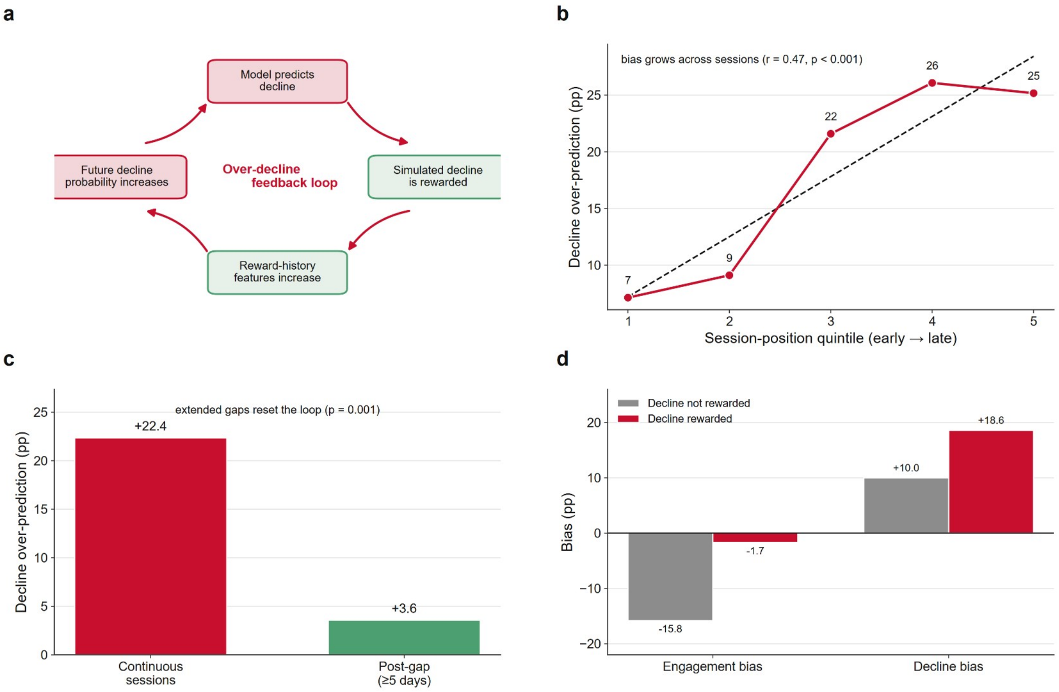


**Figure S12. Archived closed-loop simulation outputs (exploratory generative-boundary analysis).** Free-running simulation in which the model's own predictions are fed back as ongoing history. This analysis is exploratory and configuration-sensitive and is not used as primary evidence (see Supplementary Information S7); all panels are drawn from the archived Phase-3F simulation tables. (a) Candidate autoregressive over-decline feedback loop: a predicted decline is rewarded, reward-history features rise, and future decline probability increases. (b) Decline over-prediction across session-position quintiles. (c) Reset of the over-prediction after extended between-session gaps. (d) Archived Phase-3F output: effect of the reward pathway (decline rewarded versus not rewarded) on engagement and decline bias. Across seeds the archived over-decline spans approximately +16.5 to +18.6 pp with no collapsed runs. These archived values are provenance-qualified historical outputs and should not be interpreted as a single reproducible effect-size estimate.
