## Supplementary Data PDF for "History-structured forecasting of rewarded give-up behavior in a rodent metacognition task"

Bin Yin, Ya-Xin Wang, Chongyi Liu, Liya Fu
School of Psychology, Fujian Normal University, Fuzhou, China

This document reproduces the Supplementary Data items (SD1–SD9) that support the manuscript. The **complete SD1–SD9 files** are openly available in the Zenodo v1.18 deposit (DOI 10.5281/zenodo.21882292; all versions: 10.5281/zenodo.21228501; CC BY 4.0), in the `supplementary_data/` folder of the data package, alongside the full canonical dataset and model-output tables from which they derive. For readability, wide tables show a curated set of key columns here; every column is present in the archived files. Numeric values are shown to 3–4 significant figures.

---

### Supplementary Data 1. Dataset audit

Per-subject dataset-usability and parser-validation audit (24 subjects + an `ALL_SUBJECTS` summary row): session totals, usable %, validated sessions, and parser outcomes (PASS / partial-edge / multi-session / mismatch) with the effective parser-accuracy percentage. Key columns:

| subject_id | total | pct_usable | raw_only | sessions_validated | n_PASS | n_PARTIAL_EDGE | n_MISMATCH | effective_accuracy_pct |
|---|---|---|---|---|---|---|---|---|
| B 1 | 83 | 100 | 55 | 28 | 25 | 3 | 0 | 100 |
| B 2 | 87 | 100 | 60 | 27 | 23 | 3 | 0 | 100 |
| B 3 | 83 | 96.4 | 55 | 28 | 27 | 1 | 0 | 100 |
| B 4 | 83 | 94 | 56 | 27 | 24 | 2 | 0 | 100 |
| BLACK 1 | 84 | 100 | 84 | | | | | |
| BLACK 2 | 114 | 99.1 | 86 | 28 | 24 | 4 | 0 | 100 |
| BLACK 3 | 82 | 100 | 82 | | | | | |
| BLACK 4 | 116 | 96.6 | 88 | 56 | 52 | 4 | 0 | 100 |
| BLUE 1 | 85 | 100 | 85 | | | | | |
| BLUE 2 | 110 | 90 | 82 | 28 | 25 | 3 | 0 | 100 |
| BLUE 3 | 68 | 100 | 68 | | | | | |
| BLUE 4 | 113 | 97.3 | 85 | 56 | 50 | 6 | 0 | 100 |
| G 1 | 84 | 96.4 | 56 | 28 | 26 | 2 | 0 | 100 |
| G 2 | 84 | 96.4 | 57 | 27 | 22 | 4 | 0 | 100 |
| G 3 | 84 | 97.6 | 56 | 28 | 25 | 3 | 0 | 100 |
| G 4 | 84 | 89.3 | 57 | 28 | 20 | 3 | 4 | 85.71 |
| R 1 | 87 | 100 | 59 | 28 | 21 | 6 | 1 | 96.43 |
| R 2 | 83 | 100 | 56 | 27 | 22 | 3 | 1 | 96.3 |
| R 3 | 83 | 100 | 55 | 56 | 48 | 8 | 0 | 100 |
| R 4 | 83 | 98.8 | 56 | 27 | 24 | 2 | 0 | 100 |
| RED 1 | 84 | 100 | 84 | | | | | |
| RED 2 | 110 | 94.5 | 82 | 56 | 40 | 16 | 0 | 100 |
| RED 3 | 75 | 98.7 | 75 | | | | | |
| RED 4 | 46 | 100 | 46 | | | | | |
| ALL_SUBJECTS | 2095 | 97.5 | 1625 | 583 | 498 | 73 | 6 | 99 |

### Supplementary Data 2. Model comparison

Full model comparison across baseline, sequential, changepoint, hidden-Markov and recurrent architectures (model rows shown below), plus dataset key-results rows. Model families were screened on their documented eligible-trial sets, which differ across families. Model rows (key columns):

| phase | model | model_type | n_eval | action_acc | decline_AUPRC | decline_F1 | decline_ECE |
|---|---|---|---|---|---|---|---|
| 3A | Previous action | Heuristic | 82914 | 0.4284 | 0.4181 | 0.4765 | |
| 3A | History-only empirical rate | Heuristic | 82914 | 0.4356 | 0.5024 | | |
| 3B | M1 Stimulus-aware LogReg | Linear | 52570 | 0.4291 | 0.4811 | 0.421 | 0.0355 |
| 3B | M2 LightGBM sequential | Gradient boosting | 52570 | 0.4891 | 0.4934 | 0.4791 | 0.0565 |
| 3B | M5 EWM+history LightGBM | Gradient boosting | 52570 | 0.5107 | 0.5148 | 0.5096 | 0.0431 |
| 3C-1 | K=4 categorical HMM (proper) | HMM + LightGBM | 52570 | 0.486 | 0.44 | | |
| 3C-3 | B2 M5+Changepoint | Gradient boosting | 51090 | 0.5145 | 0.5378 | 0.5395 | 0.0456 |
| 3C-3 | M5+HMM_full+CP combined | HMM+CP+LightGBM | 51090 | 0.5111 | 0.532 | 0.535 | 0.0457 |
| 3D | GRU h=8 embed+CP (best) | Recurrent neural net | 39237 | 0.5188 | 0.5331 | 0.4944 | 0.0293 |
| 3D | GRU h=4 embed no-CP | Recurrent neural net | 39237 | 0.5186 | 0.517 | 0.4999 | 0.0107 |
| 3E | M5+CP temp-scaled (FINAL) | Gradient boosting | 51090 | 0.5166 | 0.5247 | 0.4243 | 0.0363 |

**Supplementary Data 2 (continued). Dataset key-results.**

Dataset key-results rows (item / value) from the same file, shown in three side-by-side blocks so the complete list stays on one page:

| item | value | item | value | item | value |
|---|---|---|---|---|---|
| canonical trial events | 213990 | K=4 cat-HMM (leaked) | 0.629 | Dedicated RT model | 2.54 |
| canonical trial events | 67 | K=4 cat-HMM (proper) | 0.440 | Dedicated RT model | 0.59 |
| session summary | 2162 | K=4 cat-HMM (proper) | 0.486 | Stored Phase 3E RT | 1.433 |
| late discrimination trials | 162304 | Disengaged state dwell | 928 | Stored Phase 3E RT | 2.662 |
| Subjects | 24 | State x Action RT variance | 0.163 | Stored Phase 3E RT | 0.808 |
| Patched sessions (repaired) | 74 | State-only RT variance | 0.0013 | RT improvement vs M5 | -31.8 |
| Patched trials added | 9844 | State stability (ARI) | 0.542 | c_t profile clusters | 3 |
| Prev action baseline | 0.4284 | M5+Changepoint (B2) | 0.5145 | Cross-context all test | 0.6266 |
| History-only empirical | 0.4356 | M5+Changepoint (B2) | 1.0488 | First-day-new-context | 0.6655 |
| History-only empirical | 0.5024 | M5+Changepoint (B2) | 0.5378 | Free-running simulation (Script 07 reimplementation, exploratory) | +17.13 |
| M1 stimulus-aware LogReg | 0.4291 | M5+Changepoint (B2) | 0.5395 | Free-running simulation (Script 07 reimplementation, exploratory) | -2.32 |
| M1 stimulus-aware LogReg | 0.4811 | M5+Changepoint (B2) | 0.0456 | Free-running simulation (Script 07 reimplementation, exploratory) | 0.4058 |
| M2 LightGBM sequential | 0.4891 | PELT changepoints | 131 | Free-running simulation (Script 07 reimplementation, exploratory) | 0/115 |
| M2 LightGBM sequential | 0.4934 | Strategy reversals (HMM) | 702 | Session drift correlation | 0.467 |
| M5 EWM history LightGBM | 0.5107 | Best GRU (h8 embed+CP) | 0.5188 | Early test decline bias | +8.1 |
| M5 EWM history LightGBM | 1.0997 | Best GRU (h8 embed+CP) | 1.1316 | Late test decline bias | +30.9 |
| M5 EWM history LightGBM | 0.5148 | Best GRU (h8 embed+CP) | 0.5331 | Post-gap decline bias | +3.6 |
| M5 EWM history LightGBM | 0.5096 | GRU RT prediction | 7.398 | Non-postgap decline bias | +22.4 |
| M5 EWM history LightGBM | 0.0431 | M5+CP temp-scaled (FINAL) | 0.5166 | Sim reliability HIGH | 2/23 |
| M5 EWM history LightGBM | 2.103 | M5+CP temp-scaled (FINAL) | 1.098 | Sim reliability MEDIUM | 4/23 |
| Early history ablation | -0.046 | M5+CP temp-scaled (FINAL) | 0.5247 | Sim reliability LOW | 17/23 |
| GLMM: decline_rate_5_z | 7.917 | M5+CP temp-scaled (FINAL) | 0.4243 | win_stay_rate vs bias | +0.483 |
| GLMM: reward_rate_5_z | -6.852 | M5+CP temp-scaled (FINAL) | 0.0363 | M5+CP temp-scaled (FINAL) | 0.476 |
| GLMM: subject intercept SD | 0.377 | Temperature scaling param | 1.3474 | M5+CP temp-scaled (FINAL) | 0.1801 |
| Strategy clusters (GLMM) | 3 | Dedicated RT model | 1.26 | | |

### Supplementary Data 3. Individual structuring-state (c_t) profiles

Per-subject profiles built from the five non-tautological structuring-state features (transition rate, context-adaptation speed, win-stay rate, lose-shift rate, gap sensitivity), with the strategy-cluster label. 23 labeled subjects:

| subject_id | transition_rate | context_adaptation_speed | win_stay_rate | lose_shift_rate | gap_sensitivity | strategy_label |
|---|---|---|---|---|---|---|
| B 1 | 0.5679 | 0.2 | 0.6311 | 0.3702 | 0.1274 | High-Decline / Accuracy-Insensitive |
| B 2 | 0.5259 | 0.4 | 0.5925 | 0.3851 | 0.0197 | High-Decline / Accuracy-Insensitive |
| B 3 | 0.5825 | 0.1 | 0.5421 | 0.4157 | 0.2155 | Persistent / Reward-Resistant |
| B 4 | 0.5047 | 0.2 | 0.7276 | 0.319 | 0.09427 | High-Decline / Accuracy-Insensitive |
| BLACK 1 | 0.5138 | 0.07 | 0.6046 | 0.2759 | 0.5081 | not_modeled |
| BLACK 2 | 0.5792 | 0.15 | 0.5211 | 0.4957 | 0.1794 | Persistent / Reward-Resistant |
| BLACK 3 | 0.6041 | 0 | 0.5321 | 0.3671 | 0.2185 | not_modeled |
| BLACK 4 | 0.6158 | 0.15 | 0.5714 | 0.4355 | 0.1091 | Reward-Sensitive Engager |
| BLUE 1 | 0.5326 | 0.1 | 0.5257 | 0.3043 | 0.2018 | not_modeled |
| BLUE 2 | 0.6174 | 0.05 | 0.5041 | 0.3954 | 0.07574 | Persistent / Reward-Resistant |
| BLUE 3 | 0.6062 | 0.1 | 0.5313 | 0.2941 | 0.3062 | not_modeled |
| BLUE 4 | 0.5653 | 0.3 | 0.5722 | 0.4177 | 0.3202 | Reward-Sensitive Engager |
| G 1 | 0.5258 | 0 | 0.5831 | 0.4312 | 0.0864 | High-Decline / Accuracy-Insensitive |
| G 2 | 0.561 | 0 | 0.5775 | 0.4288 | 0.1217 | High-Decline / Accuracy-Insensitive |
| G 3 | 0.5594 | 0.1 | 0.5428 | 0.4461 | 0.05179 | High-Decline / Accuracy-Insensitive |
| G 4 | 0.5314 | 0 | 0.6493 | 0.3665 | 0.05322 | High-Decline / Accuracy-Insensitive |
| R 1 | 0.5995 | 0.1 | 0.5353 | 0.381 | 0.5326 | Persistent / Reward-Resistant |
| R 2 | 0.5942 | 0.4 | 0.5489 | 0.4019 | 0.2553 | Persistent / Reward-Resistant |
| R 3 | 0.4642 | 0.09 | 0.7347 | 0.4794 | 0 | Reward-Sensitive Engager |
| R 4 | 0.5979 | 0.2 | 0.5328 | 0.4394 | 0.4351 | Persistent / Reward-Resistant |
| RED 1 | 0.5797 | 0.4 | 0.5075 | 0.3026 | 0.1536 | not_modeled |
| RED 2 | 0.5609 | 0.25 | 0.6075 | 0.2781 | 0.1551 | Reward-Sensitive Engager |
| RED 3 | 0.6033 | 0 | 0.5183 | 0.3596 | 0 | not_modeled |

### Supplementary Data 4. Reproducibility manifest

Manifest of the released canonical dataset and model-output tables (file, row count, role). 65 entries; the public code, with its own manifest, lockfile and `verify_reproduction.py`, is released in the Zenodo code package. First entries below (the folder and file are split for readability; the full relative paths and all 65 entries are in the complete manifest):

| folder | file | row_count | role |
|---|---|---|---|
| frozen_datasets/ | canonical_trial_event_dataset.parquet | 213990.0 | Primary frozen dataset |
| frozen_datasets/ | canonical_session_summary.csv | 2162.0 | Session-level summary |
| frozen_datasets/ | late_discrimination_trial_dataset.csv | 162304.0 | Late-discrimination subset |
| frozen_datasets/ | early_phase_history_features.csv | 103.0 | Early history features |
| frozen_datasets/modeling_views/ | view_all_subjects_full_history_flagged.parquet | 2162.0 | Modeling view |
| frozen_datasets/modeling_views/ | view_core_full_history_for_c_t.parquet | 1509.0 | Modeling view |
| frozen_datasets/modeling_views/ | view_core_late_discrimination_high_confidence.parquet | 942.0 | Modeling view |
| frozen_datasets/modeling_views/ | view_include_unscheduled_sensitivity.parquet | 2162.0 | Modeling view |
| frozen_datasets/modeling_views/ | view_no_unscheduled_sensitivity.parquet | 1938.0 | Modeling view |
| frozen_datasets/modeling_views/ | view_patch_excluded_sensitivity.parquet | 2088.0 | Modeling view |
| model_outputs/phase3a/ | phase3a_baseline_results.csv | 111.0 | Baseline model results |
| model_outputs/phase3a/ | phase3a_split_plan.csv | 539.0 | Train/test split specification |
| model_outputs/phase3b/ | phase3b_subject_effects.csv | 17.0 | GLMM subject effects |
| model_outputs/phase3b/ | phase3b_subject_strategy_profile_preliminary.csv | 3.0 | Strategy cluster profiles |
| model_outputs/phase3c1/ | phase3c1_glm_hmm_results.csv | 5.0 | GLM-HMM model selection |
| model_outputs/phase3c2/ | phase3c2_model_ladder_results.csv | 7.0 | GLM-HMM model ladder |
| model_outputs/phase3c3/ | phase3c3_subject_structuring_state_profiles.csv | 23.0 | Subject HMM state profiles |
| model_outputs/phase3d1b/ | phase3d1b_early_history_ablation_full.csv | 9.0 | Early history ablation (GRU) |
| model_outputs/phase3d1b/ | phase3d1b_changepoint_input_ablation_full.csv | 5.0 | Changepoint input ablation (GRU) |
| model_outputs/phase3e/ | phase3e1_calibration_results.csv | 4.0 | Calibration comparison |
| model_outputs/phase3e/ | phase3e1_feature_parsimony_results.csv | 9.0 | Feature ablation results |
| model_outputs/phase3e/ | phase3e_model_ablation.csv | 4.0 | Model ablation on choice trials |
| model_outputs/phase3e/ | phase3e_cross_context_results.csv | 17.0 | Cross-context evaluation |
| model_outputs/phase3e/ | phase3e_first_day_new_context.csv | 37.0 | First-day-new-context evaluation |

### Supplementary Data 5. Structuring-cause validation

Eight-model taxonomy (A–H) with temporal-order controls, in three sections. (i) Aggregate model comparison:

| model | action_acc | log_loss | decline_auprc | decline_ece | temperature |
|---|---|---|---|---|---|
| A: Triggering-only | 0.5538 | 1.06 | 0.6837 | 0.09224 | 1.711 |
| B: Group-history | 0.5239 | 1.076 | 0.6748 | 0.09542 | 1.854 |
| C: Subject-static | 0.648 | 1.147 | 0.703 | 0.07413 | 1.292 |
| D: Individual-history | 0.6683 | 0.8944 | 0.8982 | 0.05304 | 1.122 |
| E: M5+CP full | 0.6759 | 0.8933 | 0.9062 | 0.05264 | 1.093 |
| D(reset): Chronological | 0.6658 | 0.8428 | 0.906 | 0.05795 | 1.218 |
| F: Time-scrambled | 0.6126 | 1.114 | 0.773 | 0.09458 | 1.217 |
| G: Session-shuffled | 0.6523 | 0.9373 | 0.86 | 0.0593 | 1.144 |
| H: Cross-subject swap | 0.604 | 1.126 | 0.7164 | 0.0797 | 1.475 |

(ii) Statistical tests: paired Wilcoxon signed-rank (two-tailed, $n = 17$) with bootstrap 95% CIs (10,000 resamples, seed 42):

| comparison | metric | mean_diff | ci_lower | ci_upper | wilcoxon_stat | wilcoxon_p |
|---|---|---|---|---|---|---|
| D vs A | action_acc | 0.1108 | 0.06135 | 0.1661 | 1 | 0.0005312 |
| D vs A | log_loss | -0.1669 | -0.2271 | -0.1105 | 3 | 7.629e-05 |
| D vs A | decline_auprc | 0.1287 | 0.07398 | 0.1861 | 0 | 1.526e-05 |
| D vs B | action_acc | 0.1424 | 0.09595 | 0.1913 | 0 | 1.526e-05 |
| D vs B | log_loss | -0.1874 | -0.2573 | -0.1169 | 7 | 0.0002899 |
| D vs B | decline_auprc | 0.1365 | 0.08107 | 0.1946 | 0 | 1.526e-05 |
| D vs C | action_acc | 0.02145 | -0.006873 | 0.04742 | 33 | 0.07033 |
| D vs C | log_loss | -0.2486 | -0.3857 | -0.1199 | 14 | 0.001678 |
| D vs C | decline_auprc | 0.1258 | 0.07037 | 0.1841 | 2 | 4.578e-05 |
| E vs D | action_acc | 0.005952 | -0.005243 | 0.01977 | 63 | 0.5477 |
| E vs D | log_loss | 0.000175 | -0.0151 | 0.01427 | 69 | 0.7467 |
| E vs D | decline_auprc | 0.00364 | 0.0006037 | 0.007509 | 39 | 0.07968 |
| D(reset) vs F | action_acc | 0.05502 | 0.02186 | 0.09116 | 21 | 0.006653 |
| D(reset) vs F | log_loss | -0.2673 | -0.4123 | -0.1324 | 9 | 0.0005035 |
| D(reset) vs F | decline_auprc | 0.1177 | 0.06214 | 0.1763 | 3 | 7.629e-05 |
| D(reset) vs G | action_acc | 0.01344 | 0.0006079 | 0.02816 | 46 | 0.1594 |
| D(reset) vs G | log_loss | -0.09193 | -0.1634 | -0.02998 | 38 | 0.07141 |
| D(reset) vs G | decline_auprc | 0.007962 | 0.0009489 | 0.01526 | 38 | 0.07141 |
| D(reset) vs H | action_acc | 0.06379 | 0.03648 | 0.09239 | 8 | 0.0003815 |
| D(reset) vs H | log_loss | -0.2804 | -0.4221 | -0.1535 | 3 | 7.629e-05 |
| D(reset) vs H | decline_auprc | 0.09858 | 0.05163 | 0.1487 | 6 | 0.0002136 |
| D vs D(reset) | action_acc | 0.001593 | -0.01592 | 0.01566 | 53 | 0.2842 |
| D vs D(reset) | log_loss | 0.05043 | 0.01385 | 0.09215 | 41 | 0.09837 |
| D vs D(reset) | decline_auprc | 0.0002835 | -0.004847 | 0.004634 | 53 | 0.2842 |

(iii) Per-subject advantage (17 core subjects; per-contrast deltas) is provided in full in the archived file.

### Supplementary Data 6. Exploratory unseen-cohort transfer (with prior target history)

Exploratory transfer probe (Supplementary Information only): a model fitted on the 17 core subjects was applied to the 6 raw-only subjects' held-out final-40% suffix, using each target's own prior history. Pooled M5+Changepoint accuracy 58.5%; the individual-history advantage over triggering-only was not significant at $n = 6$ (exact sign-flip $p = .94$). Per-subject metrics (6 subjects × 7 model variants):

| subject_id | model | action_acc | log_loss | decline_auprc |
|---|---|---|---|---|
| BLACK 1 | A: Triggering | 0.4058 | 1.055 | 0.9918 |
| BLACK 3 | A: Triggering | 0.4846 | 1.321 | 0.4283 |
| BLUE 1 | A: Triggering | 0.6332 | 1.022 | 0.3635 |
| BLUE 3 | A: Triggering | 0.4797 | 1.324 | 0.3948 |
| RED 1 | A: Triggering | 0.5623 | 1.159 | 0.3448 |
| RED 3 | A: Triggering | 0.4723 | 1.163 | 0.3209 |
| BLACK 1 | B: Group-history | 0.4058 | 1.029 | 0.9918 |
| BLACK 3 | B: Group-history | 0.4846 | 1.304 | 0.4283 |
| BLUE 1 | B: Group-history | 0.6332 | 1.03 | 0.3635 |
| BLUE 3 | B: Group-history | 0.4797 | 1.307 | 0.3948 |
| RED 1 | B: Group-history | 0.5623 | 1.155 | 0.3448 |
| RED 3 | B: Group-history | 0.4723 | 1.159 | 0.3209 |
| BLACK 1 | D: Individual-history | 0.806 | 0.4985 | 0.9938 |
| BLACK 3 | D: Individual-history | 0.5038 | 1.72 | 0.4769 |
| BLUE 1 | D: Individual-history | 0.5967 | 1.397 | 0.3165 |
| BLUE 3 | D: Individual-history | 0.4662 | 1.766 | 0.428 |
| RED 1 | D: Individual-history | 0.5169 | 1.539 | 0.3186 |
| RED 3 | D: Individual-history | 0.4484 | 1.883 | 0.3574 |
| BLACK 1 | E: M5+CP | 0.8076 | 0.4794 | 0.9928 |
| BLACK 3 | E: M5+CP | 0.4987 | 1.652 | 0.4742 |
| BLUE 1 | E: M5+CP | 0.608 | 1.368 | 0.3279 |
| BLUE 3 | E: M5+CP | 0.4514 | 1.69 | 0.4417 |
| RED 1 | E: M5+CP | 0.5143 | 1.495 | 0.3203 |
| RED 3 | E: M5+CP | 0.4753 | 1.833 | 0.3685 |
| BLACK 1 | E(reset): real | 0.7963 | 0.4646 | 0.9927 |
| BLACK 3 | E(reset): real | 0.5013 | 1.805 | 0.4505 |
| BLUE 1 | E(reset): real | 0.5967 | 1.455 | 0.3085 |
| BLUE 3 | E(reset): real | 0.4662 | 1.77 | 0.4338 |
| RED 1 | E(reset): real | 0.5299 | 1.582 | 0.3664 |
| RED 3 | E(reset): real | 0.4484 | 1.925 | 0.3644 |
| BLACK 1 | F: Time-scrambled | 0.8287 | 0.4574 | 0.9922 |
| BLACK 3 | F: Time-scrambled | 0.4987 | 1.776 | 0.4556 |
| BLUE 1 | F: Time-scrambled | 0.6231 | 1.399 | 0.3756 |
| BLUE 3 | F: Time-scrambled | 0.4608 | 1.82 | 0.4017 |
| RED 1 | F: Time-scrambled | 0.5247 | 1.627 | 0.3216 |
| RED 3 | F: Time-scrambled | 0.426 | 1.956 | 0.3469 |
| BLACK 1 | H: Cross-subject | 0.3352 | 1.307 | 0.9958 |
| BLACK 3 | H: Cross-subject | 0.2346 | 5.09 | 0.4533 |
| BLUE 1 | H: Cross-subject | 0.1106 | 6.568 | 0.3441 |
| BLUE 3 | H: Cross-subject | 0.1351 | 5.438 | 0.3647 |
| RED 1 | H: Cross-subject | 0.1481 | 5.725 | 0.3491 |
| RED 3 | H: Cross-subject | 0.1868 | 4.583 | 0.3259 |

### Supplementary Data 7. Session-count reconciliation (compressed folder)

Session-by-session reconciliation of the 2,162 (canonical session summary) vs 2,095 (usability table) difference. Files: `excluded_67_by_phase.csv`, `excluded_67_by_subject.csv`, `excluded_67_reason_summary.csv`, `excluded_67_sessions_reconciliation.csv`, `session_count_reconciliation_note.md`.

### Supplementary Data 8. All-23-subject evaluation

Leakage-safe within-subject evaluation on every labeled subject (pooled summaries + 23 per-subject rows). The model beats prefix-derived modal action for 23/23 subjects and persistence for 21/23; 22/23 exceed their own decline-prevalence AUPRC baseline. The nominal 0.25 rate is descriptive only.

| subset | n_trials | n_subjects | action_accuracy | decline_auprc | decline_base_rate | AUPRC lift | accuracy > 0.25 | AUPRC > prevalence |
|---|---|---|---|---|---|---|---|---|
| All 23 labeled subjects (pooled) | 56077 | 23 | 0.5202 | 0.5137 | 0.2948 | 0.2189 | True | True |
| 17 core-validated subjects (within all-23 rerun) | 51090 | 17 | 0.5166 | 0.5247 | 0.3031 | 0.2216 | True | True |
| 6 raw-only subjects | 4987 | 6 | 0.5574 | 0.2704 | 0.2099 | 0.0605 | True | True |
| subject: B 1 | 2941 | 1 | 0.5478 | 0.4613 | 0.3557 | 0.1056 | True | True |
| subject: B 2 | 3286 | 1 | 0.4334 | 0.6675 | 0.3366 | 0.3309 | True | True |
| subject: B 3 | 3095 | 1 | 0.5344 | 0.2545 | 0.2533 | 0.0012 | True | True |
| subject: B 4 | 2980 | 1 | 0.5124 | 0.6059 | 0.2745 | 0.3314 | True | True |
| subject: BLACK 1 | 1232 | 1 | 0.6575 | 0.2704 | 0.2321 | 0.0383 | True | True |
| subject: BLACK 2 | 2537 | 1 | 0.5479 | 0.5476 | 0.4056 | 0.142 | True | True |
| subject: BLACK 3 | 780 | 1 | 0.4782 | 0.3234 | 0.2654 | 0.058 | True | True |
| subject: BLACK 4 | 2808 | 1 | 0.5053 | 0.3782 | 0.3451 | 0.0331 | True | True |
| subject: BLUE 1 | 796 | 1 | 0.6193 | 0.2365 | 0.1595 | 0.077 | True | True |
| subject: BLUE 2 | 2325 | 1 | 0.5841 | 0.2191 | 0.234 | -0.0149 | True | False |
| subject: BLUE 3 | 740 | 1 | 0.473 | 0.3368 | 0.2432 | 0.0936 | True | True |
| subject: BLUE 4 | 2515 | 1 | 0.4783 | 0.4914 | 0.367 | 0.1244 | True | True |
| subject: G 1 | 3008 | 1 | 0.5106 | 0.6565 | 0.3145 | 0.342 | True | True |
| subject: G 2 | 3070 | 1 | 0.485 | 0.6301 | 0.3026 | 0.3275 | True | True |
| subject: G 3 | 3188 | 1 | 0.442 | 0.6195 | 0.2867 | 0.3328 | True | True |
| subject: G 4 | 3144 | 1 | 0.4736 | 0.5664 | 0.2796 | 0.2868 | True | True |
| subject: R 1 | 3292 | 1 | 0.5428 | 0.3106 | 0.3041 | 0.0065 | True | True |
| subject: R 2 | 3214 | 1 | 0.5202 | 0.4821 | 0.3681 | 0.114 | True | True |
| subject: R 3 | 4026 | 1 | 0.4958 | 0.4516 | 0.2506 | 0.201 | True | True |
| subject: R 4 | 3263 | 1 | 0.6163 | 0.2454 | 0.236 | 0.0094 | True | True |
| subject: RED 1 | 770 | 1 | 0.5532 | 0.2403 | 0.174 | 0.0663 | True | True |
| subject: RED 2 | 2398 | 1 | 0.5917 | 0.2764 | 0.2656 | 0.0108 | True | True |
| subject: RED 3 | 669 | 1 | 0.4903 | 0.2855 | 0.1689 | 0.1166 | True | True |

### Supplementary Data 9. Raw-only subject reconstruction and verification (compressed folder)

Reconstruction/verification records for the 6 raw-only subjects (raw-file inventory with event-marker counts, behavioral-plausibility audit, raw-to-trial reconstruction result, session summary, README). Files: `6_rawonly_behavioral_plausibility_audit.csv`, `6_rawonly_raw_file_inventory.csv`, `6_rawonly_raw_reconstruction_result.csv`, `6_rawonly_session_summary.csv`, `README.md`.
