## Supplementary Data ZIP for "History-structured forecasting of rewarded give-up behavior in a rodent metacognition task": README_Supplementary_Data.md

# Supplementary Data descriptions

**Manuscript:** *History-structured forecasting of rewarded give-up behavior in a rodent metacognition task*
**Authors:** Bin Yin, Ya-Xin Wang, Chongyi Liu, Liya Fu (School of Psychology, Fujian Normal University, Fuzhou, China)

## Bundle overview

Nine curated Supplementary Data items (SD1 to SD9) that provide the specific source tables behind the numerical claims in the manuscript and Supplementary Information: dataset-audit counts (SD1), the full model comparison (SD2), per-subject structuring-state (c_t) profiles (SD3), the reproducibility manifest (SD4), the eight-model structuring-cause validation with its Wilcoxon statistics and bootstrap confidence intervals (SD5), the exploratory raw-only transfer test (SD6), the session-count reconciliation (SD7), the all-23-subject evaluation (SD8), and the raw-only reconstruction/verification records (SD9).

These are focused, claim-level tables, not the complete dataset. The full canonical dataset, model-output tables, and analysis code are openly deposited on Zenodo under concept DOI 10.5281/zenodo.21228501 (CC BY 4.0 data / BSD-3-Clause code); each item below indicates its counterpart in that deposit. Formats: UTF-8 `.xlsx` for the seven tabular items, and `.zip` for the two multi-file items (SD7 and SD9). Figure-style legends for each item also appear in the Supplementary Information ("Supplementary Data" section).

## Per-item descriptions (portal Number / Title / Description)

### Supplementary Data 1
- **File:** `Supplementary_Data_1_dataset_audit.xlsx`
- **Number:** 1
- **Title:** Dataset audit
- **Description:** Per-subject dataset-usability and parser-validation audit (24 subjects plus an all-subjects summary row): session totals, usable percentage, validated sessions, parser outcomes (PASS / partial-edge / multi-session / mismatch), and effective parser accuracy.

### Supplementary Data 2
- **File:** `Supplementary_Data_2_model_comparison.xlsx`
- **Number:** 2
- **Title:** Model comparison
- **Description:** Full model comparison across baseline, sequential, changepoint, hidden-Markov and recurrent architectures, plus dataset key-results rows. Status columns distinguish fresh released-code results from archival exploratory comparators; model families use their documented eligible-trial sets and are not presented as one matched ranked benchmark.

### Supplementary Data 3
- **File:** `Supplementary_Data_3_subject_forecasts.xlsx`
- **Number:** 3
- **Title:** Individual structuring-state (c_t) profiles
- **Description:** Per-subject structuring-state (c_t) profiles for the 23 labeled subjects, from five non-tautological features (transition rate, context-adaptation speed, win-stay rate, lose-shift rate, gap sensitivity) with the strategy-cluster label.

### Supplementary Data 4
- **File:** `Supplementary_Data_4_reproducibility_manifest.xlsx`
- **Number:** 4
- **Title:** Reproducibility manifest
- **Description:** Manifest of the released canonical dataset and model-output tables (file, row count, role); 65 entries.

### Supplementary Data 5
- **File:** `Supplementary_Data_5_structuring_cause_validation.xlsx`
- **Number:** 5
- **Title:** Structuring-cause validation
- **Description:** Eight-model structuring-cause taxonomy (A to H) with temporal-order controls and a depth-matched chronological anchor: nine aggregate rows, full per-subject model metrics, and paired Wilcoxon signed-rank tests with bootstrap 95% confidence intervals.

### Supplementary Data 6
- **File:** `Supplementary_Data_6_raw_only_transfer.xlsx`
- **Number:** 6
- **Title:** Exploratory unseen-cohort transfer test (with prior target history)
- **Description:** Exploratory transfer test (Supplementary Information): a model fitted only to the 17 core subjects, evaluated on the 6 raw-only subjects' held-out final-40% suffix, using each target's own prior history. Per-subject metrics (6 subjects x 7 model variants) plus paired contrasts. Pooled M5+Changepoint accuracy 58.5%; the individual-history advantage over triggering-only was not significant at n=6 (exact sign-flip p=.94). Generated by scripts/13.

### Supplementary Data 7
- **File:** `Supplementary_Data_7_session_reconciliation.zip`
- **Number:** 7
- **Title:** Session-count reconciliation
- **Description:** Reconciliation between the 2,162 canonical sessions and the 2,095 usability-table sessions (compressed folder: per-phase, per-subject, and per-session breakdowns plus a note).

### Supplementary Data 8
- **File:** `Supplementary_Data_8_all_23_subject_evaluation.xlsx`
- **Number:** 8
- **Title:** All-23-subject evaluation
- **Description:** Corrected within-subject chronological evaluation on every labeled subject: 56,077 test trials; pooled accuracy 0.5202 and calibrated decline AUPRC 0.5137. Includes trivial baselines (persistence/modal/prevalence): model beats persistence 21/23, prefix-modal 23/23, held-out modal 18/23; 22 of 23 exceed their decline-prevalence AUPRC baseline (exception: BLUE 2).

### Supplementary Data 9
- **File:** `Supplementary_Data_9_subject_reconstruction_verification.zip`
- **Number:** 9
- **Title:** Raw-only subject reconstruction and verification
- **Description:** Reconstruction/verification records for the 6 raw-only subjects (compressed folder: raw-file inventory with event-marker counts, behavioral-plausibility audit, raw-to-trial reconstruction result, session summary, README).
